# SARS-CoV-2 Omicron boosting induces de novo B cell response in humans

**DOI:** 10.1101/2022.09.22.509040

**Authors:** Wafaa B. Alsoussi, Sameer K. Malladi, Julian Q. Zhou, Zhuoming Liu, Baoling Ying, Wooseob Kim, Aaron J. Schmitz, Tingting Lei, Stephen C. Horvath, Alexandria J. Sturtz, Katherine M. McIntire, Birk Evavold, Fangjie Han, Suzanne M. Scheaffer, Isabella F. Fox, Luis Parra-Rodriguez, Raffael Nachbagauer, Biliana Nestorova, Spyros Chalkias, Christopher W. Farnsworth, Michael K. Klebert, Iskra Pusic, Benjamin S. Strnad, William D. Middleton, Sharlene A. Teefey, Sean P.J. Whelan, Michael S. Diamond, Robert Paris, Jane A. O’Halloran, Rachel M. Presti, Jackson S. Turner, Ali H. Ellebedy

## Abstract

The primary two-dose SARS-CoV-2 mRNA vaccine series are strongly immunogenic in humans, but the emergence of highly infectious variants necessitated additional doses of these vaccines and the development of new variant-derived ones^1–4^. SARS-CoV-2 booster immunizations in humans primarily recruit pre-existing memory B cells (MBCs)^5–9^. It remains unclear, however, whether the additional doses induce germinal centre (GC) reactions where reengaged B cells can further mature and whether variant-derived vaccines can elicit responses to novel epitopes specific to such variants. Here, we show that boosting with the original SARS-CoV-2 spike vaccine (mRNA-1273) or a B.1.351/B.1.617.2 (Beta/Delta) bivalent vaccine (mRNA-1273.213) induces robust spike-specific GC B cell responses in humans. The GC response persisted for at least eight weeks, leading to significantly more mutated antigen-specific MBC and bone marrow plasma cell compartments. Interrogation of MBC-derived spike-binding monoclonal antibodies (mAbs) isolated from individuals boosted with either mRNA-1273, mRNA-1273.213, or a monovalent Omicron BA.1-based vaccine (mRNA-1273.529) revealed a striking imprinting effect by the primary vaccination series, with all mAbs (n=769) recognizing the original SARS-CoV-2 spike protein. Nonetheless, using a more targeted approach, we isolated mAbs that recognized the spike protein of the SARS-CoV-2 Omicron (BA.1) but not the original SARS-CoV-2 spike from the mRNA-1273.529 boosted individuals. The latter mAbs were less mutated and recognized novel epitopes within the spike protein, suggesting a naïve B cell origin. Thus, SARS-CoV-2 boosting in humans induce robust GC B cell responses, and immunization with an antigenically distant spike can overcome the antigenic imprinting by the primary vaccination series.

## Main Text

The emergence of SARS-CoV-2 variants with increasing numbers of mutations in the spike protein (S) has decreased the effectiveness of primary series vaccinations and led to a recommendation for booster immunizations in most populations^10–16^. Multiple reports documented that booster immunizations based on the original Washington strain (WA1/2020) enhanced antibody responses to the ancestral strain as well as emerging variants of concern^5–8, 17–19^. In addition, new vaccines based on circulating variants were made to enhance the ability of induced antibodies to combat such variants. Indeed, recent evidence indicates that a B.1.351 (Beta)-containing booster can generate higher titers of neutralizing antibodies against both B.1.351 and B.1.1.529 (Omicron) BA.1 strains of SARS-CoV-2 compared to a booster based on the original strain alone and bivalent boosters encoding the original strain and either the BA.1 or BA.5 strain induced broader neutralizing antibody responses than the constituent monovalent vaccines^20, 21^. Whether re-exposure to mRNA vaccines encoding S from the original SARS-CoV- 2 strain or variants of concern induce robust germinal center (GC) reactions that are critical for refining high-affinity and durable antibody responses has not been examined in humans. To address these questions, we conducted an immunization study of 46 healthy adults with no history of SARS-CoV-2 infection, all of whom completed a primary vaccination course with either the Pfizer-BioNTech (BNT162b2) or Moderna (mRNA-1273) SARS-CoV-2 mRNA vaccines. Recruited individuals received a booster dose of 50 μg mRNA-1273, or mRNA- 1273.213, which contains a total of 100 μg of mRNA encoding B.1.351 and B.1.617.2 (Delta) SARS-CoV-2 S proteins (**Extended Data Table 1, 2**), as a sub-study of an ongoing clinical trial (NCT04927065).

Seven of the participants received a booster immunization of mRNA-1273, and thirty-nine received mRNA-1273.213. Blood samples were collected at baseline and weeks 1, 2, 4, 8, 17, and 26 after vaccination. Five and twenty participants in the mRNA-1273 and mRNA- 1273.213 cohorts, respectively consented to collection of fine needle aspirates (FNAs) of draining axillary lymph nodes (LNs) at weeks 2 and 8. Three and eleven participants in the mRNA-1273 and mRNA-1273.213 cohorts, respectively consented to collection of bone marrow aspirates 26 weeks after their third dose (**Fig. 1a**). Circulating S-specific antibody-producing plasmablasts (PBs) were measured by enzyme-linked immune absorbent spot (ELISpot) assay. S-specific IgG- and IgA-producing PBs were detected one week after the booster immunization from all participants in the mRNA-1273 cohort. Robust circulating IgG-producing PB responses against the original WA1/2020 strain S as well as the vaccine-encoded B.1.351 and B.1.617.2 S proteins were detected in all participants in the mRNA-1273.213 cohort 1 week after immunization, with lower IgA responses detectable in most participants (**Fig. 1b, Extended Data Fig. 1a**). Plasma IgG antibody titers against S from the WA1/2020, B.1.351, B.1.617.2, and BA.1 strains were measured by multiplex bead binding assay. In both cohorts, plasma antibody binding levels increased against all strains 4 weeks after immunization and declined slightly by 17 weeks (**Extended Data Fig. 1b**).

**Figure 1.**
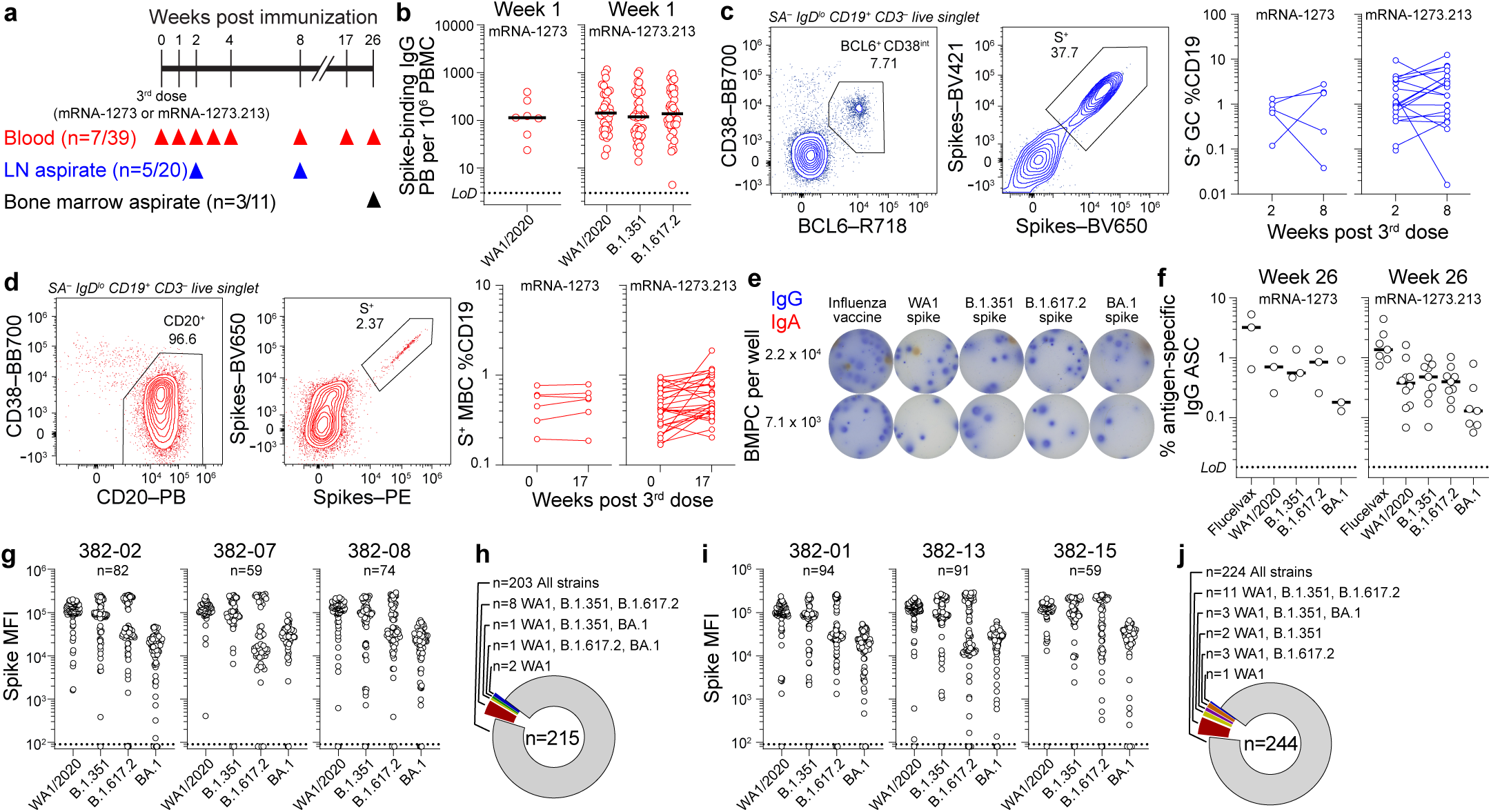
B cell response to mRNA-1273 and mRNA-1273.213 booster immunizations. (**a**) Study design. Seven and thirty-nine healthy adult volunteers were enrolled and received an mRNA-1273 or mRNA-1273.213 booster, respectively. Blood was collected at baseline and at 1, 2, 4, 8, 17, and 26 weeks post-boost. Fine needle aspirates (FNAs) of ipsilateral axillary lymph nodes were collected 2 and 8 weeks post-boost from 5 and 20 participants, and bone marrow aspirates were collected from 3 and 11 participants 26 weeks post-boost in the mRNA-1273 and mRNA-1273.213 cohorts, respectively. (**b**) Frequencies of S-binding IgG-producing plasmablasts (PB) in PBMC 1 week post-boost measured by enzyme-linked immunosorbent spot (ELISpot) in participants who received mRNA-1273 (left) and mRNA-1273.213 (right). (**c**) Representative flow cytometry plots of BCL6 and CD38 staining of streptavidin (SA)^−^ IgD^lo^ CD19^+^ CD3^−^ live singlet lymphocytes in FNA samples (left), pooled (WA1/2020, 1.351, 1.617.2, and 1.1.529 all on BV421 and BV650) S probe staining on BCL6^+^CD38^int^ GC B cells (left center), and frequencies of S^+^ GC B cells from FNA of draining lymph nodes from participants who received mRNA-1273 (right center) and mRNA-1273.213 (right). (**d**) Representative flow cytometry plots of CD20 and CD38 staining of SA^−^ IgD^lo^ CD19^+^ CD3^−^ live singlet lymphocytes in PBMC (left), pooled S probe staining on CD20^+^CD38^lo/int^ B cells (left center), and frequencies of S^+^ memory B cells (MBCs) from PBMC 17 weeks post-boost in participants who received mRNA-1273 (right center) and mRNA-1273.213 (right). (**e**) Representative ELISpot wells coated with the indicated antigens, and developed in blue (IgG) and red (IgA) after plating the indicated numbers of bone marrow plasma cells (BMPCs). (**f**) Frequencies of IgG-secreting BMPCs specific for the indicated antigens 26 weeks post-boost in participants who received mRNA-1273 (left) and mRNA-1273.213 (right). Black lines indicate medians. Symbols at each time point represent one sample. For mRNA-1273 and mRNA- 1273.213 respectively, n = 7 and 38 (b), n = 5 and 20 (c), n = 6 and 28 (d), n = 3 and 10 (f). (**g, i**) Binding of mAbs from S^+^ MBCs 17 weeks post-boost from participants who received mRNA- 1273 (g) and mRNA-1273.213 (i) to indicated strains of SARS-CoV-2 S measured by multiplex bead binding array. (**h, j**) Summary of mAb cross-reactivity from participants who received mRNA-1273 (h) and mRNA-1273.213 (j).

Ultrasonography was used to identify and guide FNA of accessible axillary nodes on the side of immunization 2 and 8 weeks post-boost. FNA samples were stained with pooled fluorescently labeled S probes from the WA1/2020, B.1.351, B.1.617.2, and BA.1 strains to detect S-specific B cells and analyzed by flow cytometry. We included BA.1 probes in the pool to detect MBCs that may have stochastically mutated to better recognize the variant antigen^22^. S- binding GC B cells, defined as CD19^+^ CD3^−^ IgD^lo^ Bcl6^+^ CD38^int^ lymphocytes, and T follicular helper cells (Tfh), defined as CD3^+^ CD19^−^ CD4^+^ CD8^−^ CD14^−^ CXCR5^+^ PD-1^+^ BCL6^+^ FoxP3^−^ lymphocytes, were detected in FNAs from all participants analysed at week 2. Frequencies of S^+^ GC B cells and Tfh correlated significantly and remained readily detectable in all but one participant from each cohort at week 8, consistent with the robust GC response observed after primary vaccination with SARS-CoV-2 mRNA vaccines (**Fig. 1c, Extended Data Fig. 1c– e**)^2, 23, 24^. Circulating MBCs in blood were identified as CD19^+^ CD3^−^ IgD^lo^ CD20^+^ lymphocytes that bound the pooled fluorescently labeled S probes. S^+^ MBCs were detected in all participants prior to boosting and at similar frequencies 17 weeks post-boost, with median frequencies of 0.52% and 0.55% of total circulating B cells in the mRNA-1273 cohort and 0.42% and 0.49% in the mRNA-1273.213 cohort (**Fig. 1d**). Consistent with sustained plasma antibody titers, bone marrow plasma cells (BMPCs) producing IgG that bound WA1/2020, B.1.351, B.1.617.2, and BA.1 S were detected in all thirteen participants with enough BMPCs for ELISpot (3 from the mRNA-1273 and 10 from the mRNA-1273.213 cohort). Frequencies of WA1/2020, B.1.351, and B.1.617.2-binding BMPCs were approximately two-fold lower than those of seasonal influenza vaccine-specific BMPCs, which accumulate over a lifetime of repeated antigen exposures. Frequencies of BA.1-binding BMPCs were approximately four-fold lower than those binding the other strains. S-binding IgA-producing BMPCs were considerably rarer or undetectable (**Fig. 1e, f, Extended Data Fig. 1f, g**).

To characterize the breadth of the MBC repertoire after boosting, we selected three participants from each cohort for whom we had characterized the B cell response to the primary vaccination series, 382-02, 382-07, and 382-08 in the mRNA-1273 cohort and 382-01, 382-13, and 382-15 in the mRNA-1273.213 cohort^2, 24^. For both cohorts, we stained week 17 post-boost PBMCs with pooled fluorescently labeled S probes from the WA1/2020, B.1.351, B.1.617.2, and BA.1 strains, allowing us to sort MBCs for mAb generation regardless of the S probe they bound. We also bulk-sorted week 17 total MBCs for heavy chain Ig sequencing to broaden the clonal repertoire analyses (**Extended Data Fig. 2a**). We generated 82, 59, 74, 94, 91, and 59 clonally distinct antigen-specific mAbs from participants 382-02, 382-07, 382-08, 382-01, 382- 13, and 382-15, respectively. We next assessed their binding to S from the WA1/2020, 1.351, 1.617.2, and BA.1 strains by multiplex bead binding assay. Confirmatory ELISAs were performed for mAbs that did not bind in the multiplex assay. Remarkably, 203 of 215 mAbs (94%) and 224 of 244 mAbs (92%) derived from the MBCs from the mRNA-1273 and mRNA- 1273.213 cohorts respectively, recognized the original WA1/2020 SARS-CoV-2 S as well as B.1.351, B.1.617.2, and BA.1 S proteins (**Fig. 1g–j, Extended Data Fig. 2b**).

**Figure 2.**
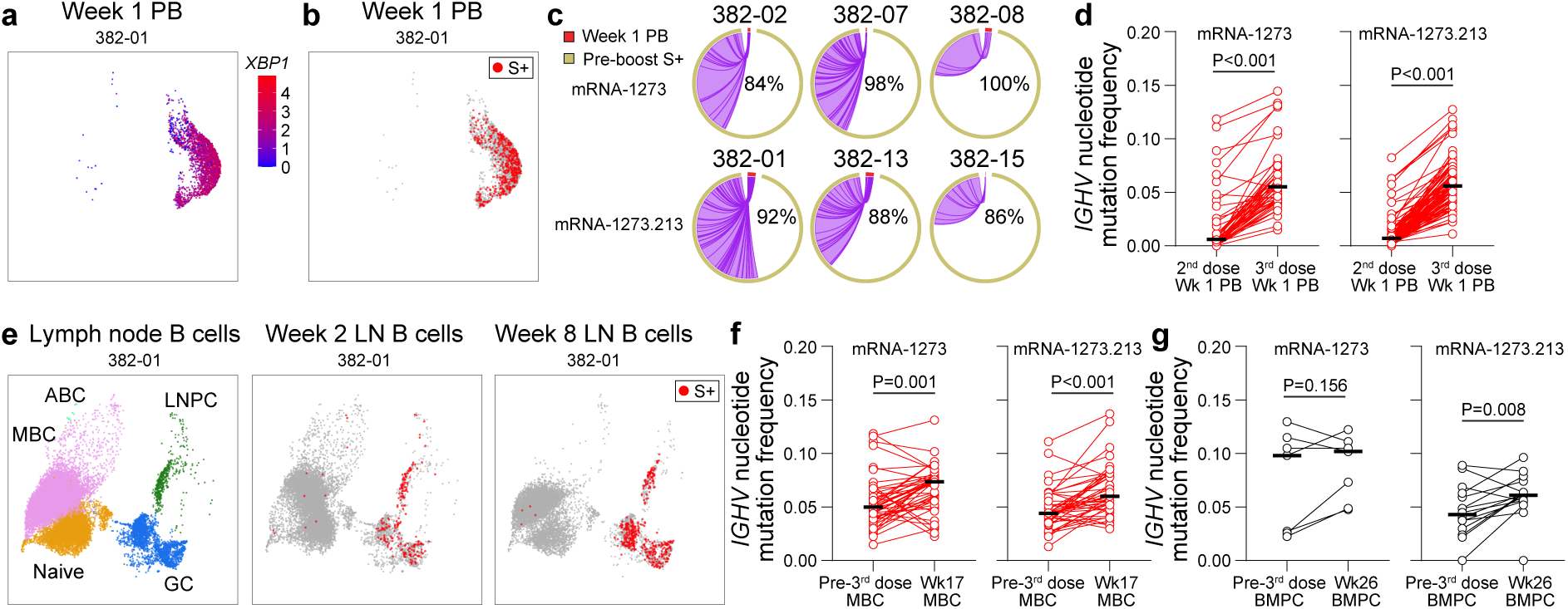
Maturation of S+ MBCs and BMPCs in response to mRNA-1273 or 1273.213 booster immunizations. (**a**, **b**, **e**) Uniform manifold approximation and projection (UMAP) of scRNA-seq transcriptional clusters of B cells either from sorted circulating PBs 1 week post boost with log-normalised *XBP1* gene expression (a) and S-specific clones (b) overlaid, or from FNAs of draining lymph nodes with S-specific clones overlaid (e). Each dot represents a cell. (**c**) Clonal overlap and percentages of S-specific PB clones related to clones generated during the primary vaccine response among participants receiving mRNA-1273 (upper) and mRNA- 1273.213 (lower). Arc length corresponds to the number of B cell receptor sequences and chord width corresponds to clone size. Purple chords correspond to overlapping clones. Percentages are of PB clones related to pre-boost S-specific clones. (**d**, **f**, **g**) Paired median immunoglobulin heavy chain variable region gene (*IGHV*) mutation frequencies of S-specific clones found in PB both 1 week after the 2^nd^ dose of the primary vaccine series and boost (d), MBCs identified both 6 months after primary vaccination and 17 weeks after boost (f), and BMPCs identified both 6 and/or 9 months after primary vaccination and 6 months after boost (g). Each symbol represents the median mutation frequency of a clone; horizontal lines indicate medians. For mRNA-1273 and mRNA-1273.213 respectively, n = 52 and 104 (d), n = 44 and 41 (f), n = 7 and 16 (g). *P* values from two-sided Wilcoxon test.

To track the clonal dynamics of the B cell response in both cohorts, we performed single-cell RNA sequencing (scRNA-seq) on week 1 sorted PB and week 2 and 8 FNA specimens from the same six participants (**Fig. 2a, e, Extended Data Fig. 3a–g**). We then linked the B cell receptor sequences to known S-specific clones identified from either week 17 MBC-derived mAbs or the previously characterized response to the primary vaccination series (**Fig. 2b, e, Extended Data Fig. 3h**)^2, 24^. The majority of S-specific PBs identified after boosting by scRNA-seq were clonally related to MBCs, GC B cells, and/or plasma cells induced by primary vaccination (**Fig. 2c**). Multiple S-specific clones were detected in the PB response after both the 2^nd^ dose of the primary vaccination series and the booster. Representatives of these clones participating in the booster PB response had significantly higher somatic hypermutation (SHM) frequencies in their immunoglobulin heavy chain variable region (IGHV) genes than those from the primary response, consistent with their recall from affinity-matured MBCs (**Fig. 2d**). S- specific PB clones identified one week post-boost were identified in GC responses in all 6 participants analyzed, though peak frequencies of the S-specific GC clonal repertoire occupied by PB clones varied widely among participants from 17% to 100% (**Extended Data Fig. 3i, j**). SHM frequencies among S-specific MBCs 17 weeks after boost were significantly higher than their clonally related counterparts isolated 6 months after primary vaccination, consistent with additional rounds of GC-driven maturation. Similar trends were observed among paired S- specific BMPC clones analyzed 6 or 9 months after primary immunization and 6 months after boost. However, SHM did not increase in all clonal families, consistent with durable populations of MBCs and BMPCs generated by the primary vaccine response persisting through the booster response (**Fig. 2f, g**). Both mRNA-1273 and mRNA-1273.213 elicited robust GC responses and maturation of the MBC and BMPC responses, but remarkably no antibodies were isolated that specifically targeted the variant strains encoded by the mRNA-1273.213 vaccine and did not cross-react to the original WA1/2020 S protein. Even among GC B cells and MBC from FNA 8 weeks after vaccination, where S-binding cells are much more frequent than in PBMC, frequencies of cells that bound the B.1.351 and B.1.617.2 strains S but not WA1/2020 were too low to sort for analysis (**Extended Data Fig. 3k, l**). Thus, the B cell response after boosting with the mRNA1273.213 vaccine was dominantly imprinted by the primary vaccination series with mRNA-1273 encoding the ancestral spike.

**Figure 3.**
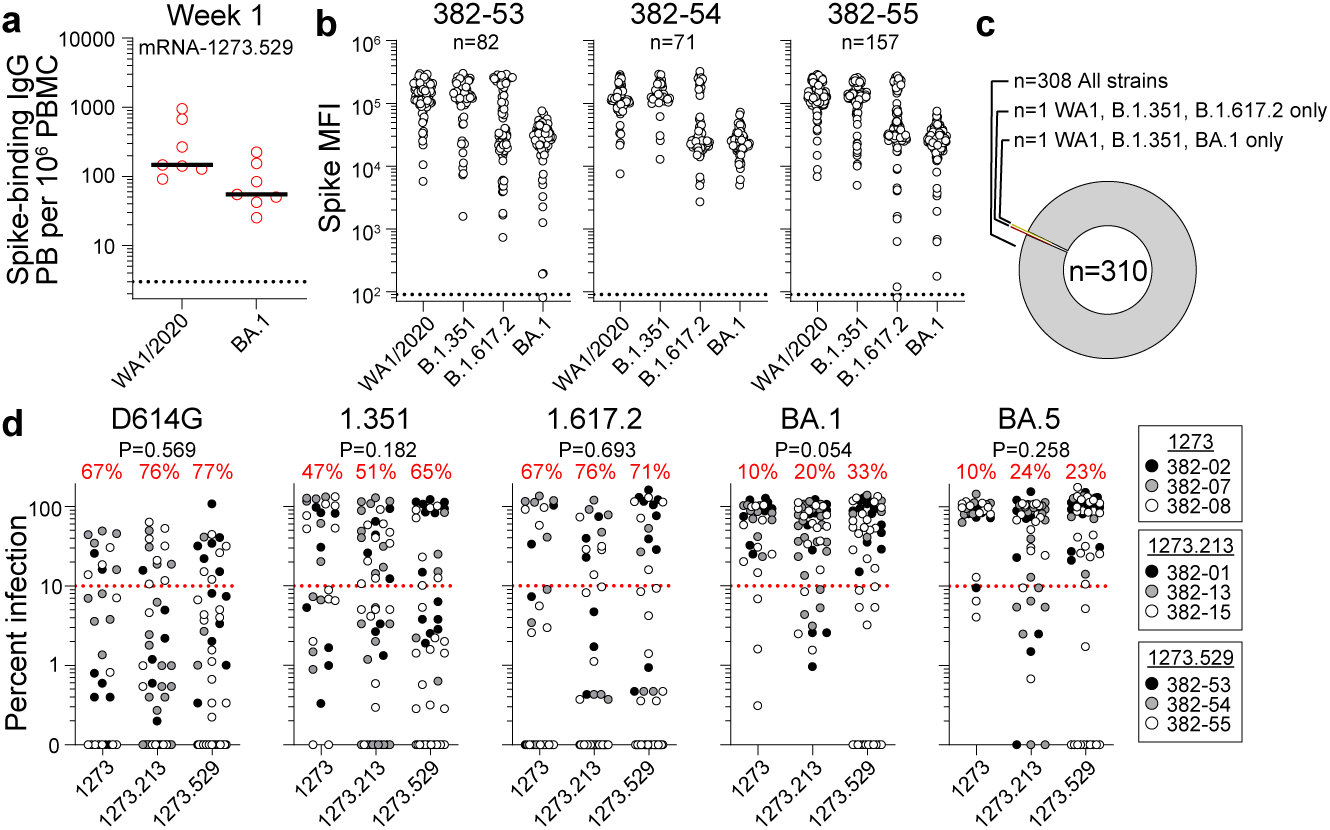
Neutralization capacity of MBC-derived mAbs. (**a**) Frequencies of S-binding IgG- producing plasmablasts (PB) in PBMC 1 week post-boost measured by ELISpot in participants who received mRNA-1273.529. Horizontal lines indicate medians. Each symbol represents 1 sample, n = 7. (**b**) Binding of mAbs from S^+^ MBCs 17 weeks post-boost to indicated strains of SARS-CoV-2 S measured by multiplex bead binding array. (**c**) Summary of mAb cross- reactivity. (**d**) Neutralizing activity of mAbs from week 17 S^+^ MBCs against indicated strains of authentic SARS-CoV-2 virus from participants who received indicated booster vaccines. Each symbol represents an individual mAb, n = 39 for mRNA-1273, n = 49 for mRNA-1273.213, and n = 52 for mRNA-1273.529. Percentages are of mAbs below the 90% infection reduction threshold. *P* values from chi-squared tests between vaccine cohorts.

To determine whether a more antigenically divergent booster could generate a detectable response targeting novel epitopes, we recruited 8 participants who had received a two-dose mRNA primary vaccination series and had no history of SARS-CoV-2 infection to receive 50 μg mRNA-1273.529, which encodes BA.1 S protein as a first or second booster after mRNA-1273 (**Extended Data Tables 1, 2**). We analysed peripheral blood samples 1 and 17 weeks post-boost with mRNA-1273.529. All 7 participants analysed at 1 week had robust circulating IgG-producing PB responses against the original WA1/2020 strain S as well as the vaccine-encoded BA.1 S protein, with lower IgA responses detectable in most participants (**Fig. 3a, Extended Data Fig. 4a**). To analyse the breadth of the MBC repertoire, we sorted MBCs from week 17 post-boost PBMCs from participants 382-53, 382-54, and 382-55 (all of whom received mRNA- 1273.529 as a first boost) using the same pooled fluorescent S probes from the WA1/2020, B.1.351, B.1.617.2, and BA.1 strains to detect MBCs regardless of their specificity and to maintain consistency with the previously generated mAbs. Like the mAbs from the mRNA-1273 and mRNA-1273.213 cohorts, 308 out of 310 mAbs (99%) were cross-reactive, binding S from WA1/2020, B.1.351, B.1.617.2, and BA.1 (**Fig. 3b, c**). To assess the neutralization capacity of the MBC-derived mAbs, we first screened all 769 mAbs from all three cohorts with a high-throughput assay employing chimeric vesicular stomatitis virus (VSV) in which the native envelope glycoprotein was replaced with S from the WA1/2020 strain with substitution D614G^25^. Thirty, 49, and 52 mAbs from the mRNA-1273, mRNA-1273.213, and mRNA- 1273.529 cohorts, respectively neutralized infection by at least 80% at 5 μg/mL (**Extended Data Fig. 4b**). We then evaluated the neutralizing capacity of these 131 mAbs against a panel of authentic, infectious SARS-CoV-2 variants, including WA1/2020 D614G, B.1.351, B.1.617.2, BA.1, and BA.5, against which 67%, 47%, 67%, 10%, and 10% from the mRNA-1273 cohort, 76%, 51%, 76%, 20%, and 24% from the mRNA-1273.213 cohort, and 77%, 65%, 71%, 33%, and 23% from the mRNA-1273.529 cohort, respectively reduced infection by at least 90% at 5 μg/mL (**Fig. 3d**).

**Figure 4.**
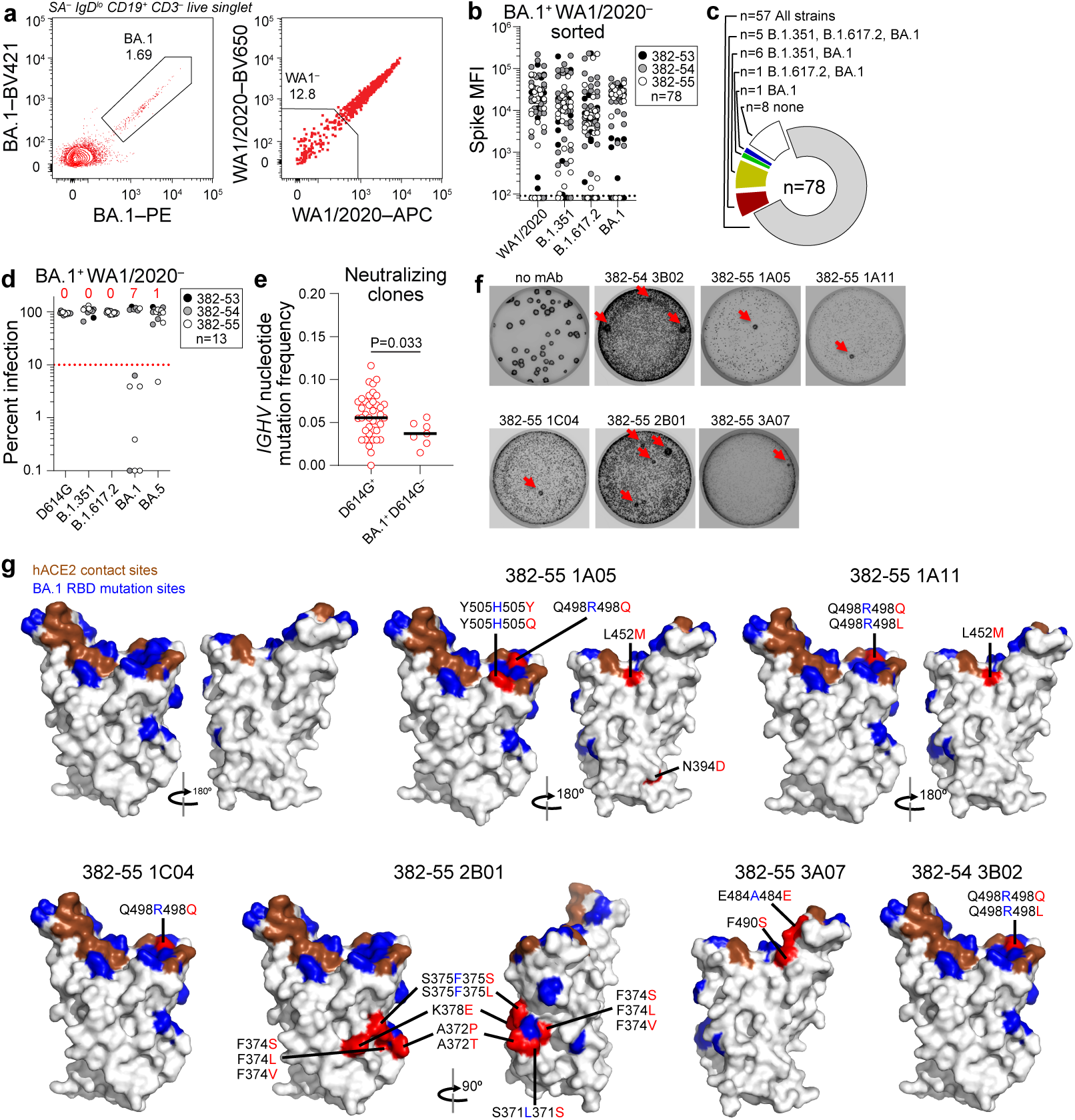
Characterization of BA.1-specific mAbs. (**a**) Gating strategy for sorting BA.1^+^ WA1/2020^−^ MBC from 17 weeks post-boost PBMC. (**b**) Binding of mAbs from BA.1^+^ WA1/2020^−^ sorted MBCs to indicated strains of SARS-CoV-2 S measured by multiplex bead binding array. (**c**) Summary of mAb binding. (**d**) Neutralizing activity of BA.1^+^ WA1/2020^−^ binding mAbs against indicated strains of authentic SARS-CoV-2 virus. Numbers above each virus are of mAbs below the 90% infection reduction threshold. (**e**) IGHV mutation frequencies of clones related to mAbs from participants 382-54 and 382-55 that neutralized D164G (left) and BA.1 but not D614G (right). Black lines indicate medians. Each symbol represents a sequence; n = 39 for D614G^+^ and n = 7 for BA.1^+^ D614G^−^. (**f**) Plaque assays on Vero E6 cells with indicated mAb in the overlay to isolate escape mutants (red arrows). Images are representative of three experiments per mAb. (**g**) Structure of RBD with hACE2 footprint highlighted in brown, BA.1 mutations highlighted in blue, and amino acids whose substitution confers resistance to indicated mAbs in plaque assays highlighted in red.

Given the high frequency of cross-reactive mAbs, we took a more targeted approach to sort MBCs from participants specific for BA.1 that did not bind WA1/2020382-53, 382-54, and 382-55 (**Fig. 4a**). Remarkably, 57 of the 78 mAbs (73%) isolated using this approach still cross-reacted with WA1/2020, indicating that the S-dependent sorting approach is not ideally sensitive for detecting low-affinity clones. Eight of the mAbs did not bind any of the antigens above background, 1 bound BA.1 only, and the remaining 12 bound BA.1 and B.1.351 and/or B.1.617.2 (**Fig. 4b, c**). We tested the neutralizing capacity of the 13 mAbs that bound BA.1 but not WA1/2020 against the panel of SARS-CoV-2 variants. As expected, none of the 13 neutralized the ancestral D614G strain. Seven of the 13 (54%) neutralized BA.1, one neutralized BA.5 (8%), and none neutralized B.1.351 or B.1.617.2 at 5 μg/mL (**Fig. 4d**). All 7 of the BA.1- neutralizing mAbs were from participants 382-54 and 382-55. We compared the SHM frequency among the clonal families of these mAbs to those from the same participants that neutralized D614G and found that the former had significantly lower levels of SHM (**Fig. 4e**). All 7 of the BA.1-neutralizing mAbs targeted the receptor binding domain (RBD) of S (**Extended Data Fig. 4c**). To define the amino acid residues targeted by the 6 most potently neutralizing BA.1-specific mAbs, we used VSV-SARS-CoV-2-S chimeric viruses (S from BA.1 strain) to select variants that escape neutralization as previously described^26, 27^. We performed plaque assays on Vero cells with the 6 neutralizing mAbs in the overlay, purified the neutralization-resistant plaques, and sequenced the S genes (**Fig. 4f, Extended Data Fig. 4d**). Sequence analysis identified S escape mutations at residue R498 for mAbs 382-54 3B02, 382-55 1A11, and 382-55 1C04; residues R498 and H505 for mAb 382-55 1A05; residues L371, A372, F374, F375, and K378 for mAb 382-55 2B01; and residue A484 for mAb 382-55 3A07. Notably, many of the mutations were reversions to the ancestral strain of SARS-CoV-2. (**Fig. 4g**).

This study evaluated antigen-specific B cell responses to SARS-CoV-2 mRNA-based booster (third or fourth) immunizations in humans. We show that boosting with an original SARS-CoV-2 S or bivalent B.1.351/B.1.617.2-matched vaccines induce robust S-specific GC response in draining axillary LNs of all sampled participants that lasted at least eight weeks after vaccination. Spike-specific PBs and GC B cells predominantly originate from pre-existing clonal lineages, which is consistent with the fact that most MBC-derived mAbs we isolated recognized the original SARS-CoV-2 S protein. We also demonstrate that immunization with the monovalent B.1.1.529-matched vaccine can induce *de novo* B cell responses against novel epitopes in the B.1.1.529 S protein. These observations expand the large body of data showing improved potency and breadth of serum antibody responses after SARS-CoV-2 booster immunization in humans^5–9, 18^.

Several critical insights relevant to cellular immunity to SARS-CoV-2 and recall responses to vaccination can be drawn from this study. B cell clones comprising the plasmablast compartment induced after the booster immunization were significantly more mutated than the same clones detected one week after completion of the primary vaccination series, a clear manifestation of the robust maturation process triggered by the primary vaccination^2, 24^.

Consistent with our previous work on B cell responses to seasonal influenza virus vaccination in humans^29, 30^, the data presented here show that pre-existing MBCs can be efficiently re-engaged into recall GC reactions. The frequencies of IgG-secreting BMPCs specific to the original SARS-CoV-2 S protein observed in this study are several fold higher than those measured seven months after mild SARS-CoV-2 infections or six months after the primary mRNA vaccination series^2, 24^. These increased frequencies are likely the result of the persistent GC responses induced after the primary vaccination series and the new GC reactions seeded by the booster immunization, highlighting the critical contribution of repeated antigen exposures to increasing antigen-specific BMPC frequency^30, 31^.

An important and surprising finding in our study is the exceptionally high percentage of circulating MBCs that recognize the S protein from the original SARS-CoV-2 strain in the individuals boosted with variant antigens, particularly as these boosters did not encode the original strain S protein. It is important to note that none of the participants from whom the mAbs were derived had documented SARS-CoV-2 infection or seroconverted against the virus nucleocapsid protein for the duration of the study. These data are consistent with MBCs generated by the primary vaccination series dominating the recall response induced by the booster and potentially out-competing clones specific for novel epitopes. It is possible that we could not detect more naïve B cell-derived mAbs specific for novel epitopes on the B.1.351, B.1.617.2, or B.1.1.529 S proteins because their affinity for the probes was below the limit of detection. The high antigenic similarity between the variant-derived S antigens and that of the original SARS-CoV-2 strain in the case of the bivalent B.1.351/B.1.617.2 vaccine may have contributed to the low frequency of *de novo* clones recognizing the former. We also speculate that an additional immunization with a variant-based vaccine may be needed to amplify the exclusively variant-specific B cell clones, similar to what has been observed upon H5N1 influenza virus immunization in humans^32^. Importantly, we note that many of the mutations selected when we cultured recombinant vesicular stomatitis virus expressing SARS-CoV-2 Omicron (BA.1) S in the presence of the *de novo* mAbs were reverted to the residues in the original strain. This suggests that newly escaped viruses are likely sensitive to potently neutralizing antibodies, including some clinically approved therapeutic ones, that were thought to be no longer useful because of the changes at the E484 residue, for example^34–38^.

The high prevalence of MBCs recognizing the original SARS-CoV-2 S protein is evidence of antigenic imprinting, in which B cell responses to previously encountered antigens remain dominant even after exposure to different but antigenically related antigens^38, 39^. The current study provides direct evidence that immunization with a SARS-CoV-2 S antigen that is sufficiently distant antigenically from the original strain can engage naïve B cells that target novel epitopes on the immunizing antigen and thus overcome such imprinting.

## Acknowledgements

We thank all the donors for generously providing precious specimens. We thank Lisa Kessels and the Washington University School of Medicine 382 Study Team (study coordinators Alem Haile, Ryley Thompson, Delaney Carani, RN, Kim Gray, MSN, APRN-BC, and Chapelle Ayres; pharmacists Michael Royal, RPh and John Tran; and laboratory technicians Laura Blair, Anita Afghanzada, and Natalie Schodl) for assistance with scheduling participants and sample collection. We thank Pamela Woodard, Betsy Thomas, Mike Harrod, Rosemary Hamlin, Maggie Rohn, and the staff of the Center for Clinical Research Imaging at Washington University School of Medicine for assistance with sample collection. We thank Claire Dalton and Brittany Roemmich for performing the nucleocapsid binding assay. The WU382 study was reviewed and approved by the Washington University Institutional Review Board (approval no. 202109021).

## Author contributions

AHE, JAO, RMP, RP, BN, and SC conceived designed the study. JAO, MKK, and RMP wrote and maintained the IRB protocol, recruited participants, and coordinated sample collection. WBA, WK, FH, and JST processed specimens. WBA, SKM, and JST performed multiplex bead array and ELISA. WBA, FH, and JST performed ELISpot. WBA and SKM performed VSV neutralization assays. WBA, SKM, WK, AJSchmitz, TL, SCH, AJSturtz, KMM, BE, IFF, and JST generated and characterized monoclonal antibodies. JQZ analyzed scRNA-seq and BCR repertoire data. ZL rescued and produced the chimeric vesicular stomatitis viruses for neutralization assays and performed and analyzed epitope mapping. BY performed the SARS- CoV-2 virus neutralization assays. WK and AJSturtz prepared libraries for scRNA-seq.

AJSchmitz performed RNA extractions and library preparation for BCR bulk sequencing and expressed SARS-CoV-2 S and variant proteins. SMS generated the authentic SARS-CoV-2 virus stocks. CWF supervised the nucleocapsid binding assay. IP supervised bone marrow specimen collection. BSS and WDM performed FNA. BSS, WDM, and SAT supervised lymph node evaluation prior to FNA and specimen collection and evaluated lymph node ultrasound data. JST sorted cells and collected and analysed the flow cytometry data. LP-R, RP, RN, JST, and AHE analyzed data. AHE, MSD, and SPJW supervised experiments and obtained funding. JST and AHE composed the manuscript. All authors reviewed and edited the manuscript.

## Data and materials availability

Antibody sequences are deposited on GenBank under the following accession numbers: xx–xx, available from GenBank/EMBL/DDBJ. Bulk sequencing reads are deposited on Sequence Read Archive under BioProject xx. The IMGT/V-QUEST database is accessible at http://www.imgt.org/IMGT_vquest/. Materials are available upon request, through a simple interinstitutional materials transfer agreement.

## Materials and Methods

### Sample collection, preparation, and storage

All studies were approved by the Institutional Review Board of Washington University in St. Louis. Written consent was obtained from all participants. Fifty-four healthy volunteers were enrolled, of whom 26 and 15 provided axillary LN and bone marrow aspirate samples, respectively (**Extended Data Table 1**). Blood samples were collected in ethylenediaminetetraacetic acid (EDTA) evacuated tubes (BD), and peripheral blood mononuclear cells (PBMC) were enriched by density gradient centrifugation over Lymphopure (BioLegend). The residual red blood cells were lysed with ammonium chloride lysis buffer, washed with PBS supplemented with 2% FBS and 2 mM EDTA (P2), and PBMC were immediately used or cryopreserved in 10% dimethylsulfoxide (DMSO) in FBS. Ultrasound-guided FNA of axillary LNs was performed by a radiologist. LN dimensions and cortical thickness were measured, and the presence and degree of cortical vascularity and location of the LN relative to the axillary vein were determined prior to each FNA. For each FNA sample, six passes were made under continuous real-time ultrasound guidance using 22- or 25-gauge needles, each of which was flushed with 3 mL of RPMI 1640 supplemented with 10% FBS and 100 U/mL penicillin/streptomycin, followed by three 1-mL rinses. Red blood cells were lysed with ammonium chloride buffer (Lonza), washed with P2, and immediately used or cryopreserved in 10% DMSO in FBS. Participants reported no adverse effects from phlebotomies or serial FNAs. Bone marrow aspirates of approximately 30 mL were collected in EDTA tubes from the iliac crest. Bone marrow mononuclear cells were enriched by density gradient centrifugation over Ficoll-Paque PLUS (Cytiva), and remaining red blood cells were lysed with ammonium chloride buffer (Lonza) and washed with P2. Bone marrow plasma cells (BMPC) were enriched from bone marrow mononuclear cells using the EasySep Human CD138 Positive Selection Kit II (StemCell Technologies) and immediately used for ELISpot or cryopreserved in 10% DMSO in FBS.

### Cell lines

Expi293F cells were cultured in Expi293 Expression Medium (Gibco). Vero- TMPRSS2 cells^40^ (a gift from Siyuan Ding, Washington University School of Medicine) were cultured at 37 °C in Dulbecco’s modified Eagle medium (DMEM) supplemented with 10% fetal bovine serum (FBS), 10 mM HEPES (pH 7.3), 1 mM sodium pyruvate, 1× nonessential amino acids, 100 U/mL of penicillin–streptomycin, and 5 μg/mL of blasticidin.

### Antigens

Recombinant soluble spike protein (S) from WA1/2020, B.1.351, B.1.617.2, B.1.1.529 (BA.1) strains of SARS-CoV-2 and their Avi-tagged counterparts were expressed as previously described ^24, 41^. Briefly, mammalian cell codon-optimized nucleotide sequences coding for the soluble ectodomain of S (GenBank: MN908947.3, amino acids 1-1213) including a C-terminal thrombin cleavage site, T4 foldon trimerization domain, and hexahistidine tag were cloned into mammalian expression vector pCAGGS. The S sequences were modified to remove the polybasic cleavage site (RRAR to A), and two pre-fusion stabilizing proline mutations were introduced (K986P and V987P, wild type numbering). For expression of Avi-tagged variants, the CDS of pCAGGS vector containing the sequence for the relevant soluble S was modified to encode 3’ Avitag insert after the 6xHIS tag (5’-HIS tag- GGCTCCGGGCTGAACGACATCTTCGAAGCCCAGAAGATTGAGTGGCATGAG-Stop-3’; HHHHHHGSGLNDIFEAQKIEWHE-) using inverse PCR mutagenesis as previously described^42^. Recombinant proteins were produced in Expi293F cells (ThermoFisher) by transfection with purified DNA using the ExpiFectamine 293 Transfection Kit (ThermoFisher). Supernatants from transfected cells were harvested 3 days post-transfection, and recombinant proteins were purified using Ni-NTA agarose (ThermoFisher), then buffer exchanged into phosphate buffered saline (PBS) and concentrated using Amicon Ultracel centrifugal filters (EMD Millipore). To biotinylate Avi-tagged S variants, the S-Avitag substrates were diluted to 40 μM and incubated for 1 h at 30℃ with 15 μg/mL BirA enzyme (Avidity) in 0.05 M bicine buffer at pH 8.3 supplemented with 10 mM ATP, 10 mM MgOAc, and 50 μM biotin. The protein was then concentrated/buffer exchanged with PBS using a 100 kDa Amicon Ultra centrifugal filter (MilliporeSigma).

To generate antigen probes for flow cytometry staining and sorting, trimeric BirA-biotinylated recombinant S from WA1/2020, B.1.351. B.1.617.2, or B.1.1.529 (BA.1) were incubated with a 1.04-fold molar excess of BV421-, BV650-, or PE-conjugated streptavidin (BioLegend) on ice, with three equal additions of S spaced every 15 min. Fifteen min after the third S addition, D-biotin was added in 6-fold molar excess to streptavidin to block any unoccupied biotin binding sites. SA-PE-Cy5 was blocked with a 6-fold molar excess of D-biotin and used as a background staining control. Bovine serum albumin (BSA) was biotinylated using the EZ-Link Micro NHS-PEG4-Biotinylation Kit (Thermo Fisher); excess unreacted biotin was removed using 7-kDa Zeba desalting columns (Pierce).

### ELISpot assay

Wells were coated with Flucelvax Quadrivalent 2019/2020 seasonal influenza virus vaccine (Sequiris), recombinant S from the WA1/2020, B.1.351, B.1.617.2, or BA.1 strains of SARS-CoV-2, or pooled anti-κ and anti-λ light chain antibodies (Cellular Technology Limited). Direct *ex-vivo* ELISpot assays were performed to determine the number of total, influenza vaccine-binding, or recombinant S-binding IgG- and IgA-secreting cells present in PBMC and enriched BMPC samples using IgG/IgA double-color ELISpot Kits (Cellular Technology Limited) according to the manufacturer’s instructions. ELISpot plates were analyzed using an ELISpot counter (Cellular Technology Limited).

### Fluorescent bead antigen binding assay

Recombinant biotinylated S from WA1/2020, B.1.351, B.1.617.2, and BA.1 strains of SARS-CoV-2 and biotinylated BSA were incubated for 30 min on ice with different fluorescence intensity peaks of the Streptavidin Coated Fluorescent Yellow Particle Kit (Spherotech) at 9.12 ng per μg beads. Beads were washed twice with 0.05% Tween 20 in PBS, resusupended in monoclonal antibodies diluted to 65 μg/mL or plasma samples diluted 1:80 in 0.05% Tween 20 in PBS, and incubated for 30 min on ice. Beads were washed twice with 0.05% Tween 20 in PBS, stained with IgG-APC-Fire750 (M1310G05, BioLegend, 1:100), incubated for 30 min on ice, washed twice with 0.05% Tween 20 in PBS, and resuspended in 2% FBS and 2 mM EDTA in PBS and acquired on an Aurora using SpectroFlo v2.2 (Cytek). Data were analyzed using FlowJo v10 (Treestar). Background-subtracted median fluorescence intensities were calculated for each sample by subtracting its median fluorescence intensity plus two times robust standard deviation for BSA and the median fluorescence intensity of an influenza virus hemagglutinin-specific monoclonal antibody or plasma collected prior to the SARS-CoV-2 pandemic for the respective spike variant.

### ELISA

Assays were performed in 96-well plates (MaxiSorp; Thermo) coated with 100 µL of recombinant S from WA1/2020, B.1.351, B.1.617.2, and BA.1 strains of SARS-CoV-2, N-terminal domain of BA.1, receptor binding domain of BA.1, or S2 domain of WA1/2020, or bovine serum albumin diluted to 1 μg/mL in PBS, and plates were incubated at 4°C overnight.

Plates then were blocked with 10% FBS and 0.05% Tween 20 in PBS. Purified mAbs were serially diluted in blocking buffer and added to the plates. Plates were incubated for 90 min at room temperature and then washed 3 times with 0.05% Tween 20 in PBS. Goat anti-human IgG-HRP (goat polyclonal, Jackson ImmunoResearch, 1:2,500) was diluted in blocking buffer before adding to plates and incubating for 60 min at room temperature. Plates were washed 3 times with 0.05% Tween 20 in PBS and 3 times with PBS before the addition of o-phenylenediamine dihydrochloride peroxidase substrate (Sigma-Aldrich). Reactions were stopped by the addition of 1 M hydrochloric acid. Optical density measurements were taken at 490 nm.

### Vesicular Stomatitis Virus (VSV)-SARS-CoV-2-S_Δ21_ eGFP-Reduction Assay

The S genes of SARS-CoV-2 isolate WA1/2020 (with D614G mutation) and B.1.351 were synthesized and replaced the native envelope glycoprotein of an infectious molecular clone of VSV, and resulting chimeric viruses expressing S protein from SARS-CoV-2 D614G or B.1.351 were used for GFP reduction neutralization tests as previously described^25^. Briefly, 10^3^ PFU of VSV-SARS-CoV-2-S_Δ21_ was incubated for 1 h at 37°C with recombinant mAbs diluted to 5 μg/mL. Antibody-virus complexes were added to Vero E6 cells in 96-well plates and incubated at 37°C for 7.5 h. Cells were subsequently fixed in 2% formaldehyde (Electron Microscopy Sciences) containing 10 μg/mL Hoechst 33342 nuclear stain (Invitrogen) for 45 min at room temperature, when fixative was replaced with PBS. Images were acquired with an InCell 2000 Analyzer (GE Healthcare) automated microscope using the DAPI and FITC channels to visualize nuclei and infected cells (i.e., eGFP-positive cells), respectively (4X objective, 4 fields per well, covering the entire well). Images were analyzed using the Multi Target Analysis Module of the InCell Analyzer 1000 Workstation Software (GE Healthcare). GFP-positive cells were identified in the FITC channel using the top-hat segmentation method and subsequently counted within the InCell Workstation software. The sensitivity and accuracy of GFP-positive cell number determinations were validated using a serial dilution of virus. The percent infection reduction was calculated from wells to which no antibody was added. A background number of GFP-positive cells was subtracted from each well using an average value determined from at least 4 uninfected wells.

### Focus reduction neutralization test

Each mAb was incubated at 5 μg/mL in DMEM supplemented with 2% FBS, 10 mM HEPES, and 100 U/mL penicillin/streptomycin with 10^2^ focus-forming units (FFU) of different of SARS-CoV-2 strains (WA1/2020 D614G, B.1.351, B.1.617.2, BA.1, and BA.5) for 1 h at 37°C^44^. Antibody-virus complexes were added to Vero- TMPRSS2 cell monolayers in 96-well plates and incubated at 37°C for 1 h. Subsequently, cells were overlaid with 1% (w/v) methylcellulose in MEM supplemented with 2% FBS. Plates were harvested 30 h (D614G, B.1.351, or B.1.617.2-infected) or 70 h (BA.1 or BA.5-infected) later by removing overlays and fixed with 4% PFA in PBS for 20 min at room temperature. Plates were washed and incubated with an oligoclonal pool of anti-S antibodies (SARS2-2, SARS2-11, SARS2-16, SARS2-31, SARS2-38, SARS2-57, and SARS2-71)^44^, and an additional oligoclonal pool of anti-S antibodies with extended reactivity (SARS2-08, -09, -10, -13, -14, -17, -20, -26, -27, -28, -31, -41, -42, -44, -49, -62, -64, -65, and -67)^45^ were included for staining BA.1 or BA.5 infected plates. Plates were subsequently incubated with HRP-conjugated goat anti-mouse IgG (Sigma 12-349) in PBS supplemented with 0.1% saponin and 0.1% bovine serum albumin.

SARS-CoV-2-infected cell foci were visualized using TrueBlue peroxidase substrate (KPL) and quantitated on an ImmunoSpot microanalyzer (Cellular Technologies Limited).

### Selection of mAb escape mutants in SARS-CoV-2 S

We used VSV-SARS-CoV-2-S (BA.1 variant) chimera to select for SARS-CoV-2 S variants that escape mAb neutralization as described previously^26, 27^. Antibody neutralization resistant mutants were recovered by plaque isolation. Briefly, plaque assays were performed to isolate the VSV-SARS-CoV-2 escape mutant on Vero cells with each tested mAb in the overlay. The concentration of each mAb in the overlay was determined by neutralization assays at a multiplicity of infection of 100. Escape clones were plaque-purified on Vero cells in the presence of mAbs, and plaques in agarose plugs were amplified on MA104 cells with the mAbs present in the medium. Viral supernatants were harvested upon extensive cytopathic effect and clarified of cell debris by centrifugation at 1,000 x g for 5 min. Aliquots were maintained at -80°C. Viral RNA was extracted from VSV-SARS-CoV-2 mutant viruses using RNeasy Mini kit (Qiagen), and the S gene was amplified using OneStep RT-PCR Kit (Qiagen). The mutations were identified by Sanger sequencing (Genewiz).

### Cell sorting and flow cytometry

Staining for flow cytometry analysis and sorting was performed using freshly isolated or cryo-preserved FNA or PBMC samples. For analysis, PBMC were incubated for 30 min on ice with purified CD16 (3G8, BioLegend, 1:100), CD32 (FUN-2, BioLegend, 1:100), CD64 (10.1, BioLegend, 1:100) and PD-1-BB515 (EH12.1, BD Horizon, 1:100) in 2% FBS and 2 mM EDTA in PBS (P2), washed twice, then were stained for 30 min on ice with WA1/2020, B.1.351, B.1.617.2, and BA.1 spike probes pre-conjugated to SA-BV650 and SA-PE, S_167-180_-PE-Cy7 tetramer, S_816-830_-APC tetramer ^46^, biotin-saturated SA-PE-Cy5, ICOS-SB436 (ISA-3, Invitrogen, 1:50), IgG-BV480 (goat polyclonal, Jackson ImmunoResearch, 1:100), IgA-FITC (M24A, Millipore, 1:500), CD8a-A532 (RPA-T8, Thermo, 1:100), CD38- BB700 (HIT2, BD Horizon, 1:500), CD71-BV421 (CY1G4, 1:400), CD20-Pacific Blue (2H7, 1:400), CD4-Spark Violet 538 (SK3, 1:400), CD19-BV750 (HIB19, 1:100), IgD-BV785 (IA6-2, 1:200), CXCR5-PE-Dazzle 594 (J252D4, 1:50), CD14-PerCP (HCD14, 1:50), CD27-PE-Fire810 (O323, 1:200), CCR7-Spark 685 (G043H7, 1:100), IgM-A700 (MHM-88, 1:400), CD3-APC-Fire810 (SK7, 1:50), and Zombie NIR (all BioLegend) diluted in Brilliant Staining buffer (BD Horizon). FNA samples were incubated for 30 min on ice with purified CD16 (3G8, BioLegend, 1:100), CD32 (FUN-2, BioLegend, 1:100), CD64 (10.1, BioLegend, 1:100) and PD-1-BB515 (EH12.1, BD Horizon, 1:100) in P2, washed twice, then stained for 30 min on ice with WA1/2020, 1.351, 1.617.2, and 1.1.529 spike probes pre-conjugated to SA-BV421 and SA-BV650, S_167-180_-APC tetramer, biotin-saturated SA-PE-Cy5, IgG-BV480 (goat polyclonal, Jackson ImmunoResearch, 1:100), IgA-FITC (M24A, Millipore, 1:500), CD8a-A532 (RPA-T8, Thermo, 1:100), CD38-BB700 (HIT2, BD Horizon, 1:500), CD20-Pacific Blue (2H7, 1:400), CD4-Spark Violet 538 (SK3, 1:400), IgM-BV605 (MHM-88, 1:100), CD19-BV750 (HIB19, 1:100), IgD-BV785 (IA6-2, 1:200), CXCR5-PE-Dazzle 594 (J252D4, 1:50), CD14-PerCP (HCD14, 1:50), CD71-PE-Cy7 (CY1G4, 1:400), CD27-PE-Fire810 (O323, 1:200), CD3-APC-Fire810 (SK7, 1:50), and Zombie NIR (all BioLegend) diluted in Brilliant Staining buffer (BD Horizon). Cells were washed twice with P2, fixed for 1 h at 25°C using the True Nuclear fixation kit (BioLegend), washed twice with True Nuclear Permeabilization/Wash buffer, stained with Ki-67-BV711 (Ki-67, BioLegend, 1:200), Blimp1-PE (646702, R&D, 1:100), FoxP3-Spark 685 (206D, BioLegend, 1:200), and Bcl6-R718 (K112-91, BD Horizon, 1:200) for 1 h at 25°C, and washed twice with True Nuclear Permeabilization/Wash buffer. Samples were resuspended in P2 and acquired on an Aurora using SpectroFlo v2.2 (Cytek). Flow cytometry data were analyzed using FlowJo v10 (Treestar).

For sorting PB, PBMC collected 1 week post-boost were stained for 30 min on ice with CD20-Pacific Blue (2H7, 1:400), CD71-FITC (CY1G4, 1:200), IgD-PerCP-Cy5.5 (IA6-2, 1:200), CD19-PE (HIB19, 1:200), CXCR5-PE-Dazzle 594 (J252D4, 1:50), CD38-PE-Cy7 (HIT2, 1:200), CD4-A700 (SK3, 1:400), and Zombie Aqua (all BioLegend) diluted in P2. Cells were washed twice, and PB (live singlet CD4^−^ CD19^+^ IgD^lo^ CD20^lo^ CD38^+^ CXCR5^lo^ CD71^+^) were sorted using a Bigfoot (Invitrogen) into PBS supplemented with 0.05% BSA and immediately processed for single cell RNAseq. For bulk sorting GC and LNPC, lymph node FNA samples collected 2 or 8 weeks post-boost were stained for 30 min on ice with purified CD16 (3G8, BioLegend, 1:100), CD32 (FUN-2, BioLegend, 1:100), CD64 (10.1, BioLegend, 1:100) and PD-1-BB515 (EH12.1, BD Horizon, 1:100) in P2, washed twice, then stained for 30 min on ice with CD20-Pacific Blue (2H7, 1:400), CD19-BV750 (HIB19, 1:100), IgD-PerCP- Cy5.5 (IA6-2, 1:200), CD71-PE (CY1G4, 1:400), CXCR5-PE-Dazzle 594 (J252D4, 1:50), CD38-PE-Cy7 (HIT2, 1:200), CD4-A700 (SK3, 1:400), and Zombie Aqua (all BioLegend) diluted in P2. Cells were washed twice, and total GC B cells (live singlet CD4^−^ CD19^+^ IgD^lo^ CD20^+^ CD38^int^ CXCR5^+^ CD71^+^) and LNPC (live singlet CD4^−^ CD19^+^ IgD^lo^ CD20^lo^ CD38^+^ CXCR5^lo^ CD71^+^) were sorted using a Bigfoot (Invitrogen) into Buffer RLT Plus (Qiagen) supplemented with 143 mM β-mercaptoethanol (Sigma-Aldrich) and immediately frozen on dry ice. For sorting memory B cells, PBMC collected 17 weeks post-boost were incubated for 10 min on ice with purified CD16 (3G8, BioLegend, 1:100), CD32 (FUN-2, BioLegend, 1:100), and CD64 (10.1, BioLegend, 1:100) in P2. For sorting S^+^ memory B cells, WA1/2020, B.1.351, B.1.617.2, and BA.1 spike probes pre-conjugated to SA-BV650 and SA-PE, biotin-saturated SA-PE-Cy5, CD20-Pacific Blue (2H7, 1:400), CD19-BV605 (HIB19, 1:100), IgD-BV785 (IA6-2, 1:200), CD3-FITC (HIT3a, 1:200), CD27-A700 (M-T271, 1:200), and Zombie NIR (all BioLegend) diluted in Brilliant Staining buffer (BD Horizon) were added and stained for an additional 30 min on ice. For sorting BA.1^+^ WA1/2020^−^ memory B cells, WA1/2020 probes pre-conjugated to SA-BV650 and SA-APC, BA.1 probes pre-conjugated to SA-BV421 and SA-PE, biotin-saturated SA-PE-Cy5, CD20-Pacific Blue (2H7, 1:400), IgD-BV785 (IA6-2, 1:200), CD19-FITC (HIB19, 1:100), CD27-PE-Fire810 (O323, 1:200), CD3-A700 (HIT3a, 1:100), and Zombie NIR (all BioLegend) diluted in Brilliant Staining buffer (BD Horizon) were added and stained for an additional 30 min on ice. Cells were washed twice, and pooled S-binding single memory B cells (live singlet CD3^−^ CD19^+^ IgD^lo^ SA-PE-Cy5^−^ pooled spikes double positive) or BA.1^+^ WA1/2020^−^ single memory B cells (live singlet CD3^−^ CD19^+^ IgD^lo^ SA-PE-Cy5^−^ BA.1^+^ WA1/2020^−^) were sorted using a Bigfoot (Invitrogen) into 96-well plates containing 2 µL Lysis Buffer (Clontech) supplemented with 1 U/μL RNase inhibitor (NEB), or total IgD^lo^ memory B cells were bulk sorted into Buffer RLT Plus (Qiagen) supplemented with 143 mM β- mercaptoethanol (Sigma-Aldrich) and immediately frozen on dry ice.

### Monoclonal antibody (mAb) generation

Antibodies were cloned as described previously^47^. Briefly, VH, Vκ, and Vλ genes were amplified by reverse transcription-PCR and nested PCR reactions from singly-sorted S^+^ memory B cells using primer combinations specific for IgG, IgM/A, Igκ, and Igλ from previously described primer sets^48^ and then sequenced. To generate recombinant antibodies, restriction sites were incorporated via PCR with primers to the corresponding heavy and light chain V and J genes. The amplified VH, Vκ, and Vλ genes were cloned into IgG1 and Igκ or Igλ expression vectors, respectively, as described previously^48–50^.

Heavy and light chain plasmids were co-transfected into Expi293F cells (Gibco) for expression, and antibody was purified using protein A agarose chromatography (Goldbio). Sequences were obtained from PCR reaction products and annotated using the ImMunoGeneTics (IMGT)/V-QUEST database (http://www.imgt.org/IMGT_vquest/)^51,52^. Mutation frequency was calculated by counting the number of nonsynonymous nucleotide mismatches from the germline sequence in the heavy chain variable segment leading up to the CDR3, while excluding the 5’ primer sequences that could be error-prone.

### Single-cell RNA-seq library preparation and sequencing

Sorted PB and LN FNA samples were processed using the following 10x Genomics kits: Chromium Next GEM Single Cell 5′ Kit v2 (PN-1000263); Chromium Next GEM Chip K Single Cell Kit (PN-1000286); BCR Amplification Kit (PN-1000253); Dual Index Kit TT Set A (PN-1000215). Chromium Single Cell 5′ Gene Expression Dual Index libraries and Chromium Single Cell V(D)J Dual Index libraries were prepared according to manufacturer’s instructions. Both gene expression and V(D)J libraries were sequenced on a Novaseq S4 (Illumina), targeting a median sequencing depth of 50,000 and 5,000 read pairs per cell, respectively.

### Bulk BCR library preparation and sequencing

RNA was purified from sorted PBs and memory B cells from PBMC, GC B cells and plasma cells from LN FNA (LNPC), and CD138-enriched BMPC using the RNeasy Plus Micro kit (Qiagen). Reverse transcription, unique molecular identifier (UMI) barcoding, cDNA amplification, and Illumina linker addition to B cell heavy chain transcripts were performed using the human NEBNext Immune Sequencing Kit (New England Biolabs) according to the manufacturer’s instructions. High-throughput 2x300bp paired-end sequencing was performed on the Illumina MiSeq platform with a 30% PhiX spike-in according to manufacturer’s recommendations, except for performing 325 cycles for read 1 and 275 cycles for read 2.

### Preprocessing of bulk sequencing BCR reads

Preprocessing of demultiplexed pair-end reads was performed using pRESTO v.0.6.2^53^) as previously described^54^, with the exception that sequencing errors were corrected using the UMIs as they were without additional clustering (**Table S3**). Previously preprocessed unique consensus sequences from samples corresponding to participants in the current study and previously reported in^24, 54^ were included without additional processing. Participants 382-01, 382-02, 382-07, 382-08, 382-13, and 382-15 correspond to previously reported 368-22, 368-20, 368-02, 368-04, 368-01, and 368-10, respectively^24, 54^. Previously preprocessed unique consensus sequences from samples corresponding to participants in the current study and reported in ^55^ were subset to those with at least two contributing reads and included.

### Preprocessing of 10x Genomics single-cell BCR reads

Demultiplexed pair-end FASTQ reads were preprocessed using Cell Ranger v.6.0.1 as previously described^24^ (**Extended Data Table 4**). Previously preprocessed single-cell BCR reads from samples corresponding to participants in the current study and reported in^24^ were included.

### V(D)J gene annotation and genotyping

Initial germline V(D)J gene annotation was performed on the preprocessed BCRs using IgBLAST v.1.17.1^56^ with the deduplicated version of IMGT/V-QUEST reference directory release 202113-2^51^. Further sequence-level and cell-level quality controls were performed as previously described^24^. Exceptions for mAb sequences triggering QC filters were handled on a case-by-case basis upon inspection as follows. Indels detected in 382-01 1B04 heavy chain and 1F01 light chain and 382-53 2G07 heavy chain were accepted. The CDR3 annotations from IMGT/V-QUEST for 382-08 1G04 heavy chain, 382-54 1D09 heavy chain, and 382-55 4H10 light chain were used in lieu of those from IgBLAST as the former had nucleotide lengths that were a multiple of 3 whereas the latter did not. Individualized genotypes were inferred based on sequences that passed all quality controls using TIgGER v.1.0.0^57^ and used to finalize V(D)J annotations. Sequences annotated as non-productively rearranged by IgBLAST were removed from further analysis.

B cell clonal lineages were inferred on a by-individual basis based on productively rearranged sequences as previously described^24^. Briefly, heavy chain-based clonal inference^58^ was performed by partitioning the heavy chains of bulk and single-cell BCRs based on common V and J gene annotations and CDR3 lengths, and clustering the sequences within each partition hierarchically with single linkage based on their CDR3s^59^.

Sequences within 0.15 normalized Hamming distance from each other were clustered as clones. Following clonal inference, full-length clonal consensus germline sequences were reconstructed using Change-O v.1.0.2^60^. Within each clone, duplicate IMGT-aligned V(D)J sequences from bulk sequencing were collapsed using Alakazam v1.1.0^60^ except for duplicates derived from different lymph nodes, time points, tissues, B cell compartments, isotypes, or biological replicates.

### BCR analysis

B cell compartment labels were treated as previously described^24^. Briefly, gene expression-based cluster annotation was used for single-cell BCRs; FACS-based sorting and magnetic enrichment were used for bulk BCRs, except that PB sorts from LN FNA were labelled LNPCs; post-2^nd^ dose week 2 IgD^lo^ enriched B cells from blood were labelled activated; and post-2^nd^ dose week 4 and post-3^rd^ dose (boost) week 17 IgD^lo^ enriched B cells from blood were labelled memory. For analysis involving the memory compartment, the memory sequences were restricted to those from blood. A heavy chain-based B cell clone was considered S-specific if it contained any sequence corresponding to a S-binding mAb that was previously reported^24, 54^ or from the current study. Clonal overlap between B cell compartments was visualized using circlize v.0.4.13^62^. Somatic hypermutation (SHM) frequency was calculated for each heavy chain sequence using SHazaM v.1.0.2^60^ as previously described^24^ by counting the number of nucleotide mismatches from the germline sequence in the variable segment leading up to the CDR3, while excluding the first 18 positions that could be error-prone due to the primers used for generating the mAb sequences.

### Processing of 10x Genomics single-cell 5′ gene expression data

Demultiplexed pair- end FASTQ reads were first preprocessed on a by-sample basis and samples were subsequently subsampled to the same effective sequencing length and aggregated using Cell Ranger v.6.0.1 as previously described^24^. Quality control was performed on the aggregate gene expression matrix consisting of 336,960 cells and 36,601 features using SCANPY v.1.7.2^62^. Briefly, to remove presumably lysed cells, cells with mitochondrial content greater than 17.5% of all transcripts were removed. To remove likely doublets, cells with more than 8,000 features or 80,000 total UMIs were removed. To remove cells with no detectable expression of common endogenous genes, cells with no transcript for any of a list of 34 housekeeping genes^24^ were removed. The feature matrix was subset, based on their biotypes, to protein-coding, immunoglobulin, and T cell receptor genes that were expressed in at least 0.05% of the cells in any sample. The resultant feature matrix contained 15,751 genes. Finally, cells with detectable expression of fewer than 200 genes were removed. After quality control, there were a total of 312,242 cells from 39 single-cell samples (**Extended Data Table 4**).

### Single-cell gene expression analysis

Transcriptomic data was analyzed using SCANPY v.1.7.2^62^ as previously described^24^ with minor adjustments suitable for the current data. Briefly, overall clusters were first identified using Leiden graph-clustering with resolution 0.50 (**Extended Data Fig. 3B, Extended Data Table 5**). UMAPs were faceted by participant and inspected for convergence to assess whether there was a need for integration (**Extended Data Fig. 3C**). Cluster identities were assigned by examining the expression of a set of marker genes^63^ for different cell types (**Extended Data Fig. 3D**). To remove potential contamination by platelets, 205 cells with a log-normalized expression value of >2.5 for PPBP were removed. From a cluster consisting primarily of monocytes, 36 cells originating from LN FNA and with a log-normalized expression value of >0 for at least two of FDCSP, CXCL14^64^, and FCAMR^65^ were annotated FDCs. Cells from the overall B cell cluster were further clustered to identify B cell subsets using Leiden graph-clustering resolution 0.35 (**Extended Data Fig. 3E, Extended Data Table 6**). Cluster identities were assigned by examining the expression of a set of marker genes^63^ for different B cell subsets (**Extended Data Fig. 3F**) along with the availability of BCRs. Despite being clustered with B cells during overall clustering, one group tended to have both BCRs and relatively high expression levels of CD2 and CD3E; one group tended to have no BCRs and relatively high CD2 and CD3E; and two unassigned groups tended to have no BCRs. These were excluded from the final B cell clustering. Ten cells that were found in the GC B cell cluster but came from blood were labelled ‘PB-like’^63^. 407 cells that were found in the PB cluster but came from LN FNA were re-assigned as LNPCs. One cell that was found in the LNPC cluster but came from blood was re-assigned as PB. Heavy chain SHM frequency and isotype usage of the B cell subsets were inspected for consistency with expected values to further confirm their assigned identities.

## Extended Data figure captions

**Extended Data Figure 1.**
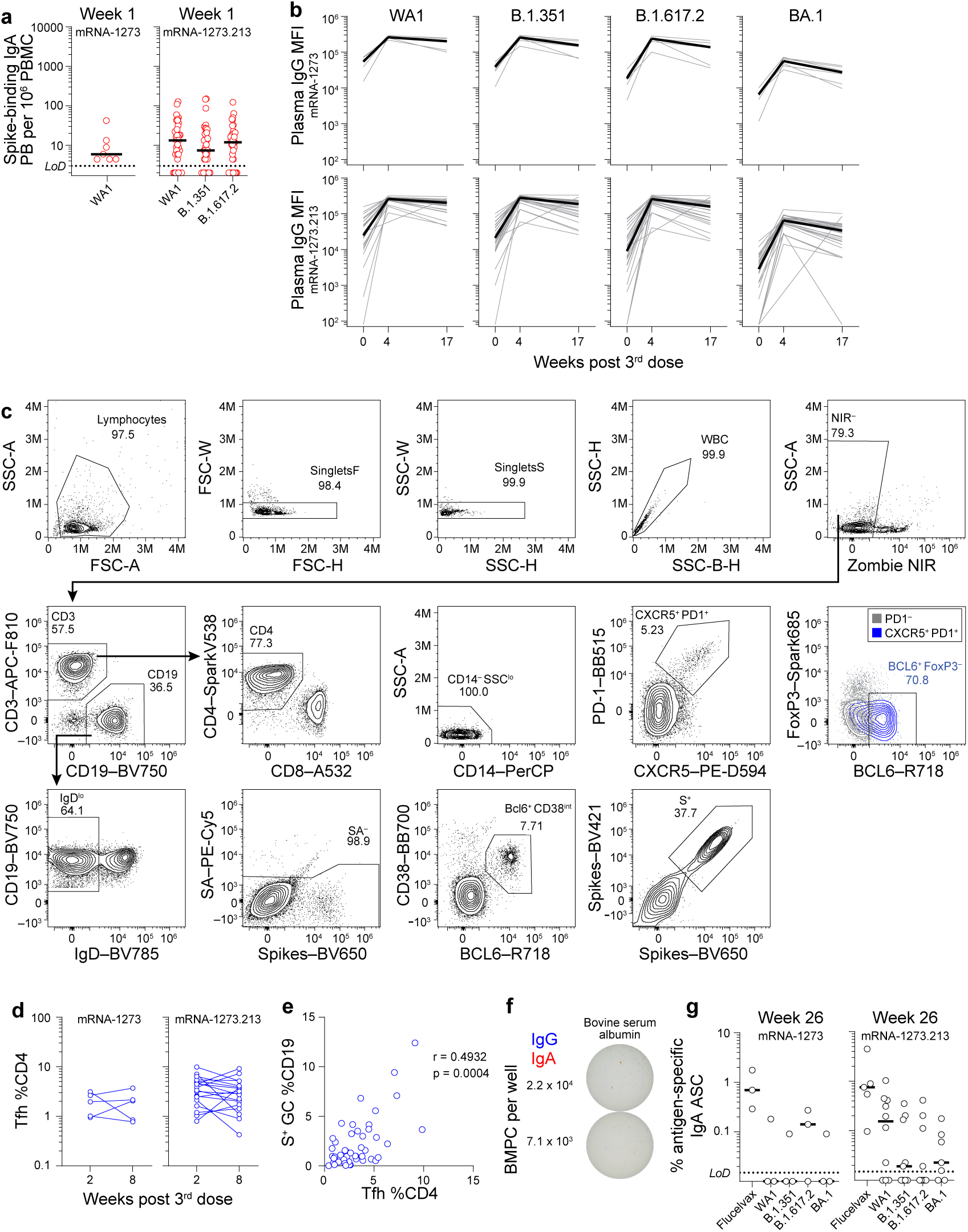
Robust GC and Tfh responses to mRNA-1273 and mRNA- 1273.213 boosters. (**a**) Frequencies of S-binding IgA-producing PB in PBMC 1 week post-boost measured by ELISpot in participants who received mRNA-1273 (left) and mRNA-1273.213 (right). (**b**) Plasma IgG binding to indicated strains of SARS-CoV-2 S measured by multiplex bead binding array in participants who received mRNA-1273 (upper) and mRNA-1273.213 (lower). (**c**) Gating strategy for analyzing S^+^ GC B cells and Tfh in FNA. (**d**) Frequencies of T follicular helper cells (Tfh) from FNA of draining lymph nodes. (**e**) Correlation between frequencies of S^+^ GC B cells and Tfh. (**f**) Representative ELISpot wells coated with BSA, and developed in blue (IgG) and red (IgA) after plating the indicated numbers of BMPCs. (**g**) Frequencies of IgA-secreting BMPCs specific for the indicated antigens 26 weeks post-boost. Black lines indicate medians. Symbols at each time point represent one sample. For mRNA-1273 and mRNA-1273.213 respectively, n = 7 and 38 (a), n = 6 and 28 (b), n = 5 and 20 (d), n = 3 and 10 (g).

**Extended Data Figure 2.**
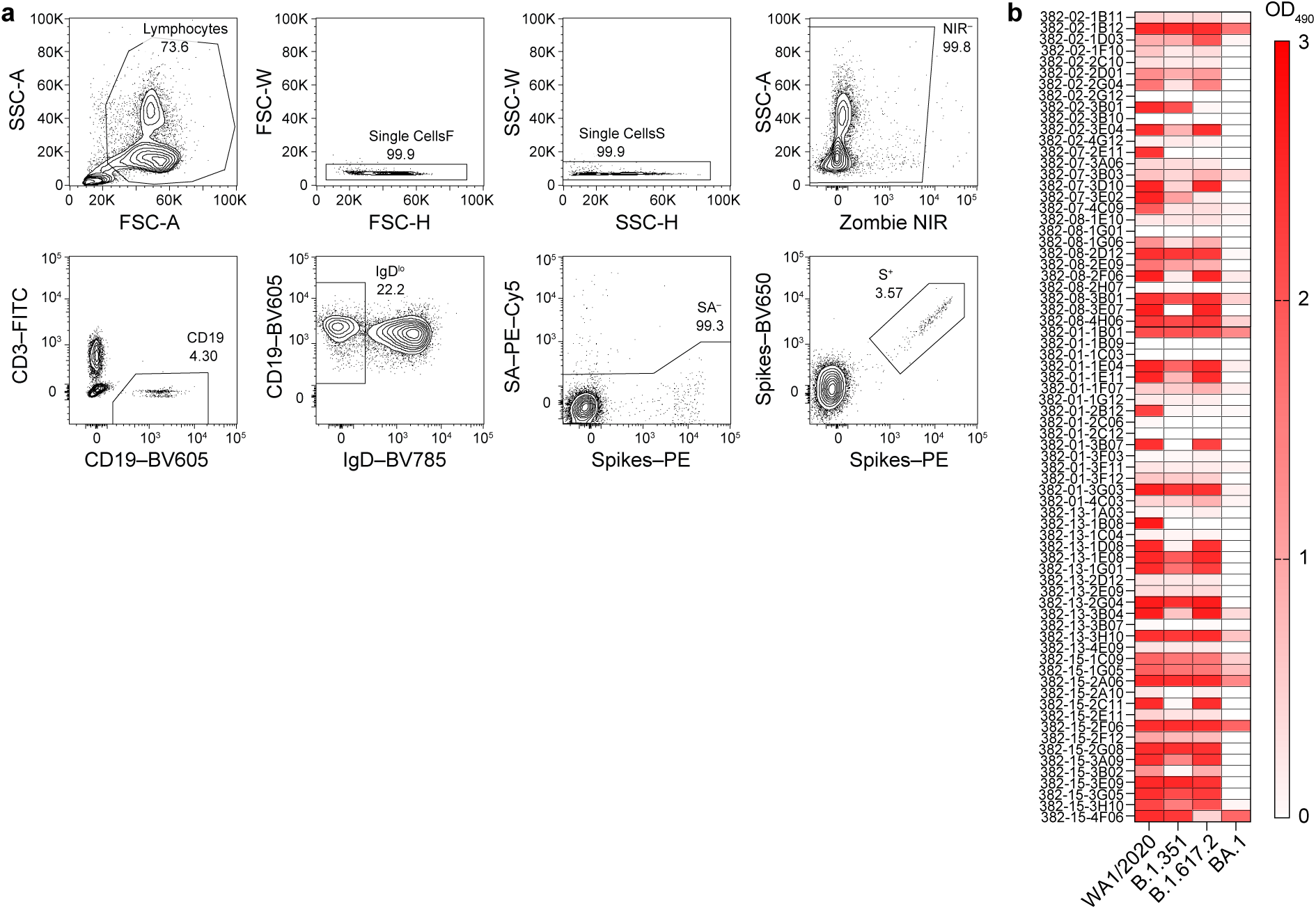
Breadth of MBC-derived mAbs after mRNA-1273 and mRNA- 1273.213 boosters. (**a**) Gating strategy for sorting S^+^ MBC from PBMC. (**b**) Binding of mAbs to indicated antigens by ELISA performed in duplicate, presented as OD_490_ minus two times the background signal to BSA.

**Extended Data Figure 3.**
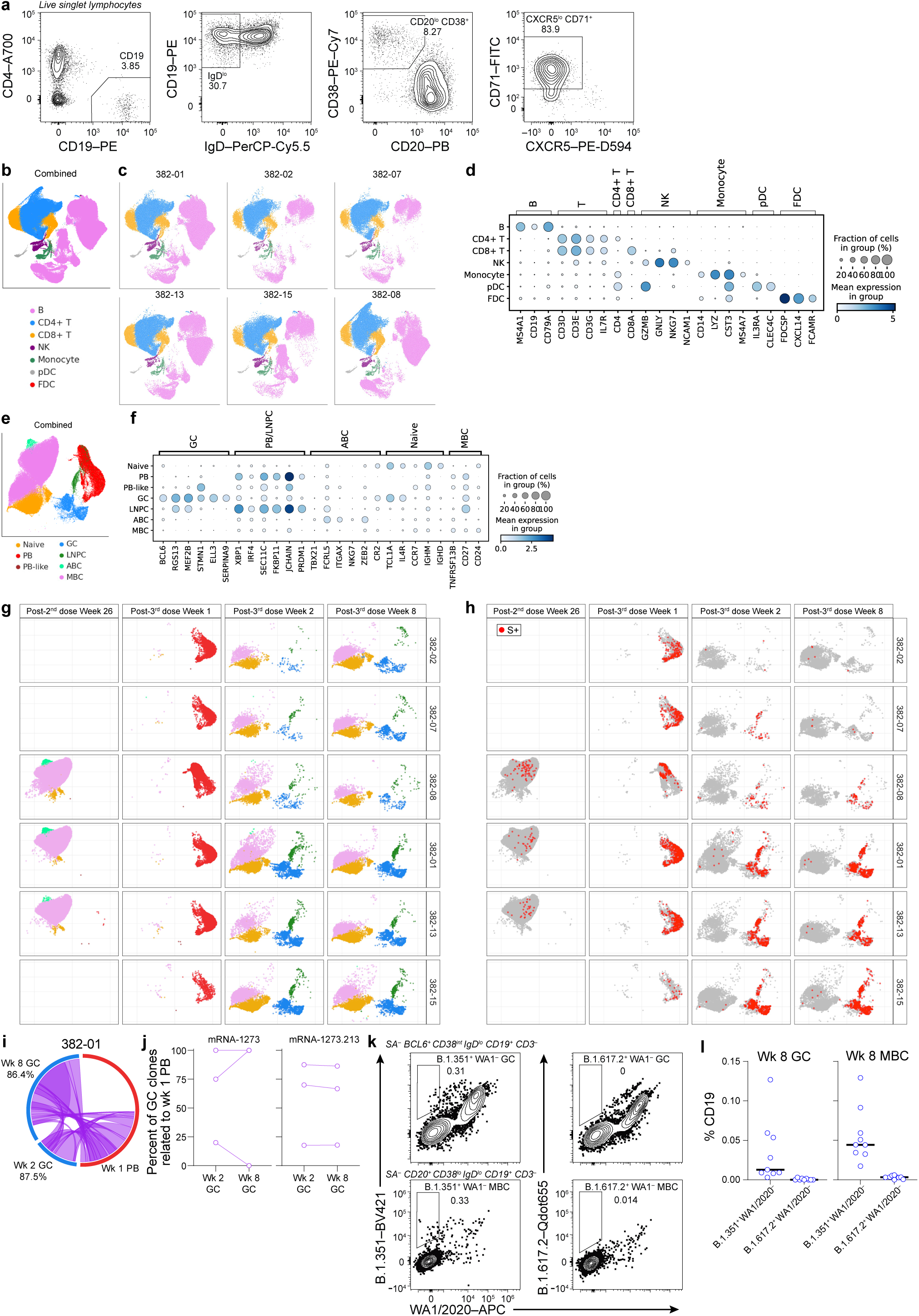
Maturation of S+ MBCs in response to mRNA-1273 or mRNA- 1273.213 booster. (**a**) Gating strategy for sorting PB from PBMC. (**b**, **c**, **e**, **g**) UMAPs showing scRNA-seq transcriptional clusters of total cells (b, c) or B cells (e, g) from all participants (b, e) or from each participant separately (c, g). (**d**, **f**) Dot plots for the marker genes used for identifying the annotated clusters in (b, c) (d) and in (e, g) (f). (**h**) SARS-CoV-2 S+ clones visualized in red on UMAPs of B cells from each participant separately and faceted by time point. (**i**) Clonal overlap between S-binding PBs and GC B cells at indicated time points. Arc length corresponds to the number of BCR sequences and chord width corresponds to clone size. Purple chords correspond to clones spanning both compartments. Percentages are of GC B cell clones related to PBs at each time point. (**j**) Percentages of S-specific GC clones related to week 1 PBs. Symbols at each time point represent one sample, n = 6. (**k**) Representative flow cytometry plots of WA1/2020 and B.1.351 (left) or B.1.617.2 (right) staining of SA^−^ BCL6^+^CD38^int^ IgD^lo^ CD19^+^ CD3^−^ live singlet lymphocytes (top) or SA^−^ CD20^+^CD38^lo^ IgD^lo^ CD19^+^ CD3^−^ live singlet lymphocytes (bottom) in FNA samples from participants who received mRNA-1273.213. (**l**) Frequencies of B.1.351^+^ WA1/2020^−^ and B.1.617.2^+^ WA1/2020^−^ GC B cells (left) and MBC (right) from FNA of draining lymph nodes from participants who received mRNA-1273.213. Black lines indicate medians. Symbols represent one sample; n = 9.

**Extended Data Figure 4.**
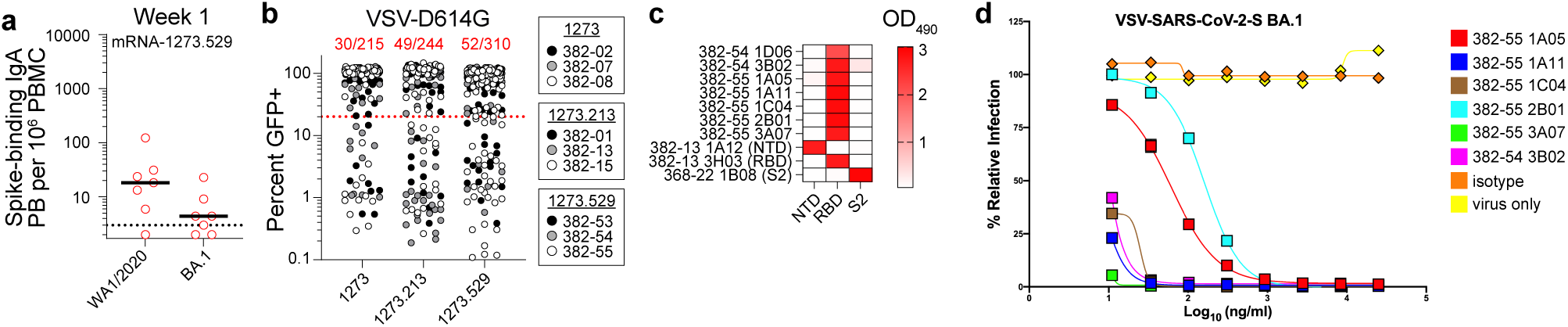
Characterization of BA.1-specific mAbs. (**a**) Frequencies of S-binding IgA-producing PB in PBMC 1 week post-boost measured by ELISpot in participants who received mRNA-1273.529. Black lines indicate medians. Symbols represent one sample; n = 7. (**b**) Neutralizing activity of mAbs from week 17 S^+^ MBCs against chimeric vesicular stomatitis virus in which the native envelope glycoprotein was replaced with S from WA1/2020 (with D614G mutation). (**c**) Binding of mAbs to BA.1 S and its constituent domains by ELISA performed in duplicate, presented as OD_490_ values. S2-specific mAb 368-22 1B08 was described previously^54^. (**d**) Titration of mAbs to determine neutralizing concentrations against VSV-SARS-CoV-2. Data are representative of two independent experiments.

## Extended Data Tables

**Extended Data Table 1.** Study WU382 participant demographics.

| **Variable** | **Total n=54**<br>**n (%)** |
| --- | --- |
| **Age (median [range])** | 37.5 (18-72) |
| **Sex** |  |
| Female | 31 (57.4) |
| Male | 23 (42.6) |
| **Race** |  |
| White | 49 (90.7) |
| Black | 2 (3.7) |
| Asian | 2 (3.7) |
| Other | 1 (1.9) |
| **Ethnicity** |  |
| Not of Hispanic, Latinx, or Spanish origin | 52 (96.3) |
| Hispanic, Latinx, Spanish origin | 2 (3.7) |
| **BMI (median [range])** | 26.86 (19.3-47.74) |
| **Comorbidities** |  |
| Lung disease | 1 (1.9) |
| Diabetes mellitus | 1 (1.9) |
| Hypertension | 10 (18.5) |
| Cardiovascular | 0 (0) |
| Liver disease | 0 (0) |
| Chronic kidney disease | 0 (0) |
| Cancer on chemotherapy | 0 (0) |
| Hematological malignancy | 0 (0) |
| Pregnancy | 0 (0) |
| Neurological | 0 (0) |
| Rheumatologic disease | 0 (0) |
| HIV | 0 (0) |
| Solid organ transplant recipient | 0 (0) |
| Bone marrow transplant recipient | 0 (0) |
| Hyperlipidemia | 2 (3.7) |
| **Type of baseline vaccine** |  |
| Pfizer | 29 (53.7) |
| Moderna | 25 (46.3) |
| **Type of booster vaccine (1<sup>st</sup> booster)** |  |
| Moderna mRNA-1273 | 12 (22.2) |
| Moderna mRNA-1273.213 | 39 (72.2) |
| Moderna mRNA-1273.529 | 3 (6) |
| **Time from second dose to booster in days**<br>**(median [range])** | 272.5 (196-356) |
| **Participants who received 2<sup>nd</sup> booster** | **5 (9)** |
| Moderna 1273.529 | 5 (9) |

**Extended Data Table 2.** Booster vaccine side effects^a^.

| **Side effects from**<br>**booster vaccines** | **Overall**<br>**(n=54)** | **mRNA-1273.213**<br>**(n=39)** | **mRNA-1273.529**<br>**(n=3)** | **mRNA-1273**<br>**(n=12)** |
| --- | --- | --- | --- | --- |
| Chills | 24 (44.4) | 21 (53.8) | 0 (0) | 2 (28.6) |
| Feeling unwell | 29 (53.7) | 25 (64.1) | 1 (33.3) | 3 (25) |
| Fever | 15 (27.8) | 14 (35.9) | 0 (0) | 1 (14.3) |
| Headache | 25 (46.3) | 20 (51.3) | 1 (33.3) | 1 (14.3) |
| Injection site pain | 40 (74.1) | 29 (74.4) | 1 (33.3) | 6 (85.7) |
| Injection site swelling | 5 (9.3) | 5 (12.8) | 0 (0) | 0 (0) |
| Injection site redness | 7 (13) | 6 (15.4) | 0 (0) | 1 (8.3) |
| Joint pain | 13 (24.1) | 10 (25.6) | 0 (0) | 3 (25) |
| Muscle pain | 24 (44.4) | 19 (48.7) | 1 (33.3) | 4 (33.3) |
| Nausea | 10 (18.5) | 8 (20.5) | 1 (33.3) | 1 (8.3) |
| Swollen lymph nodes | 14 (25.9) | 10 (25.6) | 0 (0) | 4 (25.6) |
| Tiredness | 36 (66.7) | 27 (69.2) | 1 (33.3) | 8 (66.7) |

<sup>a</sup>These data are reported for a subset of the participants enrolled in a parallel clinical trial
(NCT04927065) who provided additional consent for the collection of samples in this study.
Safety and reactogenicity data for all participants in the clinical trial will be reported separately.

**Extended Data Table 3.**
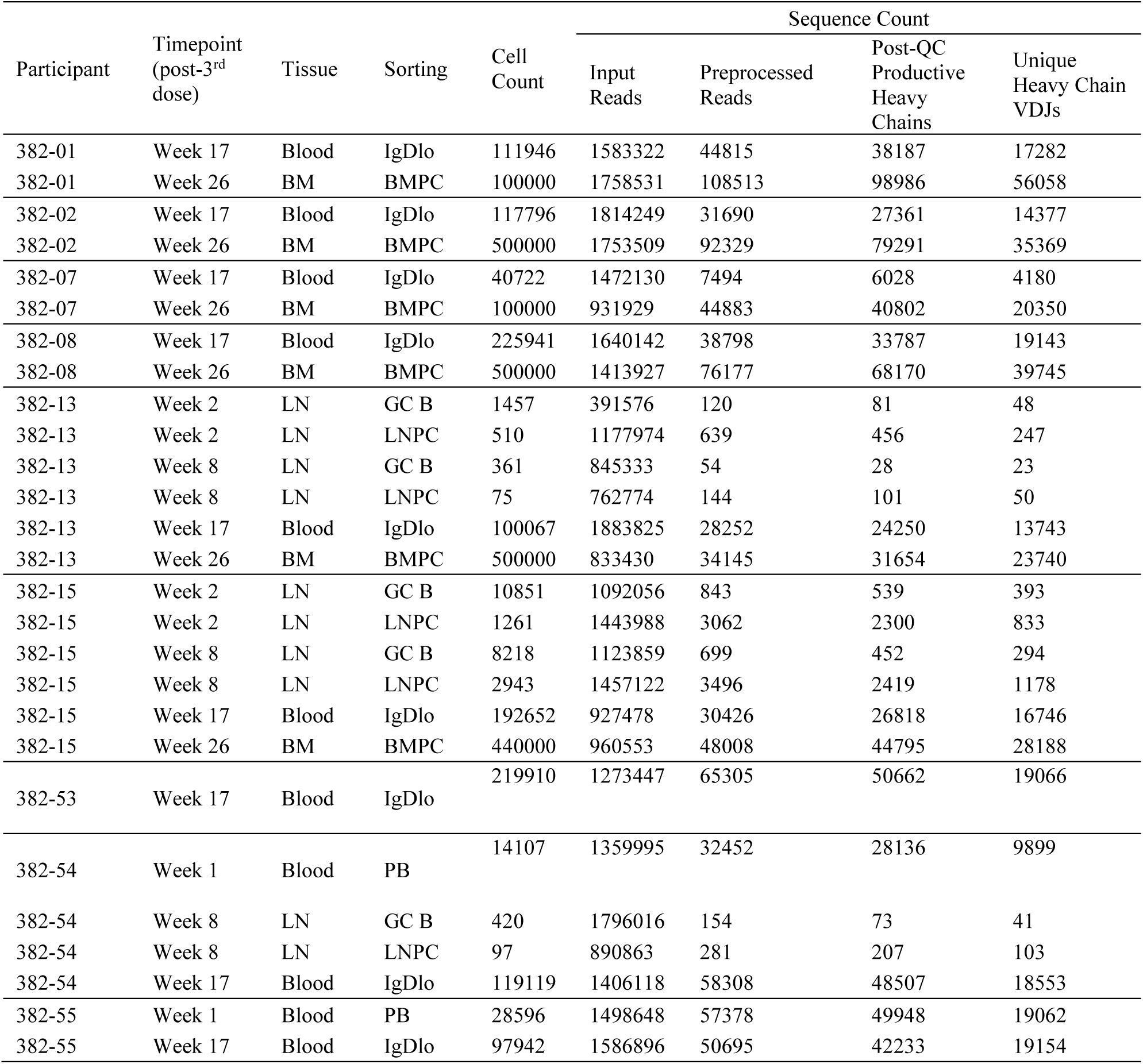
Processing of BCR reads from bulk-seq.

| Participant | Timepoint<br>(post-3<sup>rd</sup><br>dose) | Tissue | Sorting | Cell<br>Count | Sequence Count |  |  |  |
| --- | --- | --- | --- | --- | --- | --- | --- | --- |
|  |  |  |  |  | Input<br>Reads | Preprocessed<br>Reads | Post-QC<br>Productive<br>Heavy<br>Chains | Unique<br>Heavy Chain<br>VDJs |
| 382-01 | Week 17 | Blood | IgDlo | 111946 | 1583322 | 44815 | 38187 | 17282 |
| 382-01 | Week 26 | BM | BMPC | 100000 | 1758531 | 108513 | 98986 | 56058 |
| 382-02 | Week 17 | Blood | IgDlo | 117796 | 1814249 | 31690 | 27361 | 14377 |
| 382-02 | Week 26 | BM | BMPC | 500000 | 1753509 | 92329 | 79291 | 35369 |
| 382-07 | Week 17 | Blood | IgDlo | 40722 | 1472130 | 7494 | 6028 | 4180 |
| 382-07 | Week 26 | BM | BMPC | 100000 | 931929 | 44883 | 40802 | 20350 |
| 382-08 | Week 17 | Blood | IgDlo | 225941 | 1640142 | 38798 | 33787 | 19143 |
| 382-08 | Week 26 | BM | BMPC | 500000 | 1413927 | 76177 | 68170 | 39745 |
| 382-13 | Week 2 | LN | GC B | 1457 | 391576 | 120 | 81 | 48 |
| 382-13 | Week 2 | LN | LNPC | 510 | 1177974 | 639 | 456 | 247 |
| 382-13 | Week 8 | LN | GC B | 361 | 845333 | 54 | 28 | 23 |
| 382-13 | Week 8 | LN | LNPC | 75 | 762774 | 144 | 101 | 50 |
| 382-13 | Week 17 | Blood | IgDlo | 100067 | 1883825 | 28252 | 24250 | 13743 |
| 382-13 | Week 26 | BM | BMPC | 500000 | 833430 | 34145 | 31654 | 23740 |
| 382-15 | Week 2 | LN | GC B | 10851 | 1092056 | 843 | 539 | 393 |
| 382-15 | Week 2 | LN | LNPC | 1261 | 1443988 | 3062 | 2300 | 833 |
| 382-15 | Week 8 | LN | GC B | 8218 | 1123859 | 699 | 452 | 294 |
| 382-15 | Week 8 | LN | LNPC | 2943 | 1457122 | 3496 | 2419 | 1178 |
| 382-15 | Week 17 | Blood | IgDlo | 192652 | 927478 | 30426 | 26818 | 16746 |
| 382-15 | Week 26 | BM | BMPC | 440000 | 960553 | 48008 | 44795 | 28188 |
| 382-53 | Week 17 | Blood | IgDlo | 219910 | 1273447 | 65305 | 50662 | 19066 |
| 382-54 | Week 1 | Blood | PB | 14107 | 1359995 | 32452 | 28136 | 9899 |
| 382-54 | Week 8 | LN | GC B | 420 | 1796016 | 154 | 73 | 41 |
| 382-54 | Week 8 | LN | LNPC | 97 | 890863 | 281 | 207 | 103 |
| 382-54 | Week 17 | Blood | IgDlo | 119119 | 1406118 | 58308 | 48507 | 18553 |
| 382-55 | Week 1 | Blood | PB | 28596 | 1498648 | 57378 | 49948 | 19062 |
| 382-55 | Week 17 | Blood | IgDlo | 97942 | 1586896 | 50695 | 42233 | 19154 |

**Extended Data Table 4.**
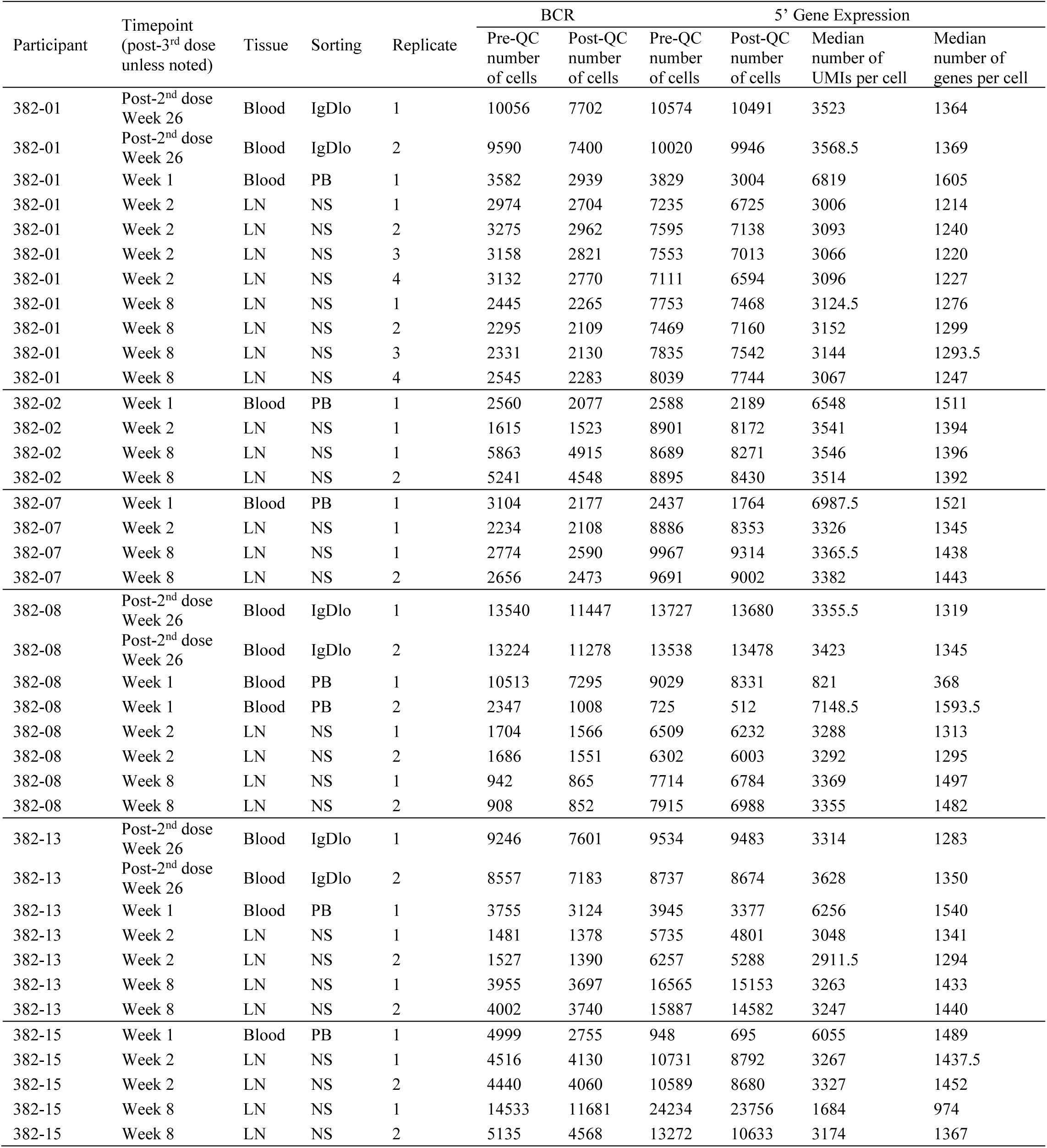
Processing of BCR and 5’ gene expression data from scRNA-seq.

| Participant | Timepoint<br>(post-3<sup>rd</sup> dose<br>unless noted) | Tissue | Sorting | Replicate | BCR |  | 5' Gene Expression |  |  |  |
| --- | --- | --- | --- | --- | --- | --- | --- | --- | --- | --- |
|  |  |  |  |  | Pre-QC<br>number<br>of cells | Post-QC<br>number<br>of cells | Pre-QC<br>number<br>of cells | Post-QC<br>number<br>of cells | Median<br>number of<br>UMIs per cell | Median<br>number of<br>genes per cell |
| 382-01 | Post-2<sup>nd</sup> dose<br>Week 26 | Blood | IgDlo | 1 | 10056 | 7702 | 10574 | 10491 | 3523 | 1364 |
| 382-01 | Post-2<sup>nd</sup> dose<br>Week 26 | Blood | IgDlo | 2 | 9590 | 7400 | 10020 | 9946 | 3568.5 | 1369 |
| 382-01 | Week 1 | Blood | PB | 1 | 3582 | 2939 | 3829 | 3004 | 6819 | 1605 |
| 382-01 | Week 2 | LN | NS | 1 | 2974 | 2704 | 7235 | 6725 | 3006 | 1214 |
| 382-01 | Week 2 | LN | NS | 2 | 3275 | 2962 | 7595 | 7138 | 3093 | 1240 |
| 382-01 | Week 2 | LN | NS | 3 | 3158 | 2821 | 7553 | 7013 | 3066 | 1220 |
| 382-01 | Week 2 | LN | NS | 4 | 3132 | 2770 | 7111 | 6594 | 3096 | 1227 |
| 382-01 | Week 8 | LN | NS | 1 | 2445 | 2265 | 7753 | 7468 | 3124.5 | 1276 |
| 382-01 | Week 8 | LN | NS | 2 | 2295 | 2109 | 7469 | 7160 | 3152 | 1299 |
| 382-01 | Week 8 | LN | NS | 3 | 2331 | 2130 | 7835 | 7542 | 3144 | 1293.5 |
| 382-01 | Week 8 | LN | NS | 4 | 2545 | 2283 | 8039 | 7744 | 3067 | 1247 |
| 382-02 | Week 1 | Blood | PB | 1 | 2560 | 2077 | 2588 | 2189 | 6548 | 1511 |
| 382-02 | Week 2 | LN | NS | 1 | 1615 | 1523 | 8901 | 8172 | 3541 | 1394 |
| 382-02 | Week 8 | LN | NS | 1 | 5863 | 4915 | 8689 | 8271 | 3546 | 1396 |
| 382-02 | Week 8 | LN | NS | 2 | 5241 | 4548 | 8895 | 8430 | 3514 | 1392 |
| 382-07 | Week 1 | Blood | PB | 1 | 3104 | 2177 | 2437 | 1764 | 6987.5 | 1521 |
| 382-07 | Week 2 | LN | NS | 1 | 2234 | 2108 | 8886 | 8353 | 3326 | 1345 |
| 382-07 | Week 8 | LN | NS | 1 | 2774 | 2590 | 9967 | 9314 | 3365.5 | 1438 |
| 382-07 | Week 8 | LN | NS | 2 | 2656 | 2473 | 9691 | 9002 | 3382 | 1443 |
| 382-08 | Post-2<sup>nd</sup> dose<br>Week 26 | Blood | IgDlo | 1 | 13540 | 11447 | 13727 | 13680 | 3355.5 | 1319 |
| 382-08 | Post-2<sup>nd</sup> dose<br>Week 26 | Blood | IgDlo | 2 | 13224 | 11278 | 13538 | 13478 | 3423 | 1345 |
| 382-08 | Week 1 | Blood | PB | 1 | 10513 | 7295 | 9029 | 8331 | 821 | 368 |
| 382-08 | Week 1 | Blood | PB | 2 | 2347 | 1008 | 725 | 512 | 7148.5 | 1593.5 |
| 382-08 | Week 2 | LN | NS | 1 | 1704 | 1566 | 6509 | 6232 | 3288 | 1313 |
| 382-08 | Week 2 | LN | NS | 2 | 1686 | 1551 | 6302 | 6003 | 3292 | 1295 |
| 382-08 | Week 8 | LN | NS | 1 | 942 | 865 | 7714 | 6784 | 3369 | 1497 |
| 382-08 | Week 8 | LN | NS | 2 | 908 | 852 | 7915 | 6988 | 3355 | 1482 |
| 382-13 | Post-2<sup>nd</sup> dose<br>Week 26 | Blood | IgDlo | 1 | 9246 | 7601 | 9534 | 9483 | 3314 | 1283 |
| 382-13 | Post-2<sup>nd</sup> dose<br>Week 26 | Blood | IgDlo | 2 | 8557 | 7183 | 8737 | 8674 | 3628 | 1350 |
| 382-13 | Week 1 | Blood | PB | 1 | 3755 | 3124 | 3945 | 3377 | 6256 | 1540 |
| 382-13 | Week 2 | LN | NS | 1 | 1481 | 1378 | 5735 | 4801 | 3048 | 1341 |
| 382-13 | Week 2 | LN | NS | 2 | 1527 | 1390 | 6257 | 5288 | 2911.5 | 1294 |
| 382-13 | Week 8 | LN | NS | 1 | 3955 | 3697 | 16565 | 15153 | 3263 | 1433 |
| 382-13 | Week 8 | LN | NS | 2 | 4002 | 3740 | 15887 | 14582 | 3247 | 1440 |
| 382-15 | Week 1 | Blood | PB | 1 | 4999 | 2755 | 948 | 695 | 6055 | 1489 |
| 382-15 | Week 2 | LN | NS | 1 | 4516 | 4130 | 10731 | 8792 | 3267 | 1437.5 |
| 382-15 | Week 2 | LN | NS | 2 | 4440 | 4060 | 10589 | 8680 | 3327 | 1452 |
| 382-15 | Week 8 | LN | NS | 1 | 14533 | 11681 | 24234 | 23756 | 1684 | 974 |
| 382-15 | Week 8 | LN | NS | 2 | 5135 | 4568 | 13272 | 10633 | 3174 | 1367 |

**Extended Data Table 5.** Cell counts and frequencies of overall transcriptional clusters from scRNA-seq.

| Participant | Transcriptional Cluster |  |  |  |  |  |  |
| --- | --- | --- | --- | --- | --- | --- | --- |
|  | B | CD4+ T | CD8+ T | NK | Monocyte | pDC | FDC |
| 382-01 | 43604 (54.1%) | 27552 (34.2%) | 6663 (8.3%) | 1114 (1.4%) | 1260 (1.6%) | 452 (0.6%) | 6 (0%) |
| 382-02 | 8722 (32.2%) | 14566 (53.9%) | 2796 (10.3%) | 249 (0.9%) | 546 (2%) | 155 (0.6%) | 11 (0%) |
| 382-07 | 8982 (31.6%) | 15493 (54.5%) | 2645 (9.3%) | 566 (2%) | 494 (1.7%) | 246 (0.9%) | 5 (0%) |
| 382-08 | 40884 (65.9%) | 15078 (24.3%) | 4685 (7.6%) | 652 (1.1%) | 432 (0.7%) | 266 (0.4%) | 1 (0%) |
| 382-13 | 31918 (52%) | 22362 (36.4%) | 5976 (9.7%) | 507 (0.8%) | 485 (0.8%) | 107 (0.2%) | 1 (0%) |
| 382-15 | 34299 (65.3%) | 13031 (24.8%) | 3466 (6.6%) | 972 (1.8%) | 633 (1.2%) | 143 (0.3%) | 12 (0%) |
| Combined | 168409 (54%) | 108082 (34.6%) | 26231 (8.4%) | 4060 (1.3%) | 3850 (1.2%) | 1369 (0.4%) | 36 (0%) |

**Extended Data Table 6.** Cell counts and frequencies of B cell transcriptional clusters from scRNA-seq.

| Participant | Timepoint<br>(post-3<sup>rd</sup> dose<br>unless noted) | B cell transcriptional cluster |  |  |  |  |  |  |
| --- | --- | --- | --- | --- | --- | --- | --- | --- |
|  |  | GC | LNPC | PB | PB-like | ABC | Naive | MBC |
| 382-01 | Post-2nd dose<br>week 26 | 0 (0%) | 0 (0%) | 0 (0%) | 1 (0%) | 470 (2.3%) | 96 (0.5%) | 19674 (97.2%) |
| 382-01 | Week 1 | 0 (0%) | 0 (0%) | 2600 (99.4%) | 0 (0%) | 0 (0%) | 3 (0.1%) | 13 (0.5%) |
| 382-01 | Week 2 | 1184 (11.1%) | 269 (2.5%) | 0 (0%) | 0 (0%) | 9 (0.1%) | 3619 (34%) | 5569 (52.3%) |
| 382-01 | Week 8 | 1228 (14.5%) | 226 (2.7%) | 0 (0%) | 0 (0%) | 0 (0%) | 1848 (21.7%) | 5195 (61.1%) |
| 382-01 | Combined | 2412 (5.7%) | 495 (1.2%) | 2600 (6.2%) | 1 (0%) | 479 (1.1%) | 5566 (13.3%) | 30451 (72.5%) |
| 382-02 | Week 1 | 0 (0%) | 0 (0%) | 1805 (99.1%) | 1 (0.1%) | 0 (0%) | 6 (0.3%) | 10 (0.5%) |
| 382-02 | Week 2 | 68 (4.9%) | 19 (1.4%) | 0 (0%) | 0 (0%) | 0 (0%) | 720 (51.6%) | 588 (42.2%) |
| 382-02 | Week 8 | 364 (7.5%) | 52 (1.1%) | 0 (0%) | 0 (0%) | 0 (0%) | 1639 (33.6%) | 2830 (57.9%) |
| 382-02 | Combined | 432 (5.3%) | 71 (0.9%) | 1805 (22.3%) | 1 (0%) | 0 (0%) | 2365 (29.2%) | 3428 (42.3%) |
| 382-07 | Week 1 | 0 (0%) | 0 (0%) | 1438 (98.9%) | 0 (0%) | 1 (0.1%) | 4 (0.3%) | 11 (0.8%) |
| 382-07 | Week 2 | 81 (4.1%) | 68 (3.5%) | 0 (0%) | 0 (0%) | 1 (0.1%) | 943 (48%) | 871 (44.3%) |
| 382-07 | Week 8 | 425 (8.8%) | 69 (1.4%) | 0 (0%) | 0 (0%) | 0 (0%) | 1403 (29.1%) | 2917 (60.6%) |
| 382-07 | Combined | 506 (6.1%) | 137 (1.7%) | 1438 (17.5%) | 0 (0%) | 2 (0%) | 2350 (28.5%) | 3799 (46.1%) |
| 382-08 | Post-2nd dose<br>week 26 | 0 (0%) | 0 (0%) | 0 (0%) | 1 (0%) | 297 (1.1%) | 246 (0.9%) | 26601 (98%) |
| 382-08 | Week 1 | 0 (0%) | 0 (0%) | 8771 (99.7%) | 1 (0%) | 0 (0%) | 0 (0%) | 29 (0.3%) |
| 382-08 | Week 2 | 157 (5.7%) | 75 (2.7%) | 0 (0%) | 0 (0%) | 3 (0.1%) | 1281 (46.6%) | 1235 (44.9%) |
| 382-08 | Week 8 | 161 (9.6%) | 57 (3.4%) | 0 (0%) | 0 (0%) | 0 (0%) | 739 (44.2%) | 715 (42.8%) |
| 382-08 | Combined | 318 (0.8%) | 132 (0.3%) | 8771 (21.7%) | 2 (0%) | 300 (0.7%) | 2266 (5.6%) | 28580 (70.8%) |
| 382-13 | Post-2nd dose<br>week 26 | 0 (0%) | 0 (0%) | 7 (0%) | 2 (0%) | 45 (0.2%) | 109 (0.6%) | 17940 (99.1%) |
| 382-13 | Week 1 | 0 (0%) | 0 (0%) | 2714 (98.7%) | 3 (0.1%) | 0 (0%) | 9 (0.3%) | 23 (0.8%) |
| 382-13 | Week 2 | 913 (31.4%) | 339 (11.7%) | 0 (0%) | 0 (0%) | 0 (0%) | 877 (30.2%) | 777 (26.7%) |
| 382-13 | Week 8 | 873 (13.6%) | 116 (1.8%) | 0 (0%) | 0 (0%) | 0 (0%) | 2478 (38.5%) | 2968 (46.1%) |
| 382-13 | Combined | 1786 (5.9%) | 455 (1.5%) | 2721 (9.0%) | 5 (0%) | 45 (0.1%) | 3473 (11.5%) | 21708 (71.9%) |
| 382-15 | Week 1 | 0 (0%) | 0 (0%) | 562 (93.4%) | 1 (0.2%) | 0 (0%) | 2 (0.3%) | 37 (6.1%) |
| 382-15 | Week 2 | 1223 (16.1%) | 173 (2.3%) | 0 (0%) | 0 (0%) | 0 (0%) | 3013 (39.8%) | 3170 (41.8%) |
| 382-15 | Week 8 | 990 (16.8%) | 298 (5.1%) | 0 (0%) | 0 (0%) | 2 (0%) | 2722 (46.2%) | 1881 (31.9%) |
| 382-15 | Combined | 2213 (15.7%) | 471 (3.3%) | 562 (4.0%) | 1 (0%) | 2 (0%) | 5737 (40.8%) | 5088 (36.2%) |
| Combined |  | 7667 (5.4%) | 1761 (1.2%) | 17897<br>(12.5%) | 10 (0%) | 828 (0.6%) | 21757<br>(15.2%) | 93054 (65.1%) |

**Extended Data Table 7.** Counts of B cells found in S-binding clones and frequencies out of respective B cell transcriptional clusters.

| Participant | Timepoint<br>(post-3<sup>rd</sup> dose<br>unless noted) | B cell transcriptional cluster |  |  |  |  |  |  |
| --- | --- | --- | --- | --- | --- | --- | --- | --- |
|  |  | GC | LNPC | PB | PB-like | ABC | Naive | MBC |
| 382-01 | Post-2nd dose<br>week 26 | 0 (-) | 0 (-) | 0 (-) | 0 (0%) | 1 (0.2%) | 0 (0%) | 31 (0.2%) |
| 382-01 | Week 1 | 0 (-) | 0 (-) | 484 (18.6%) | 0 (-) | 0 (-) | 0 (0%) | 0 (0%) |
| 382-01 | Week 2 | 125 (10.6%) | 74 (27.5%) | 0 (-) | 0 (-) | 0 (0%) | 2 (0.1%) | 9 (0.2%) |
| 382-01 | Week 8 | 315 (25.7%) | 72 (31.9%) | 0 (-) | 0 (-) | 0 (-) | 0 (0%) | 4 (0.1%) |
| 382-01 | Combined | 440 (18.2%) | 146 (29.5%) | 484 (18.6%) | 0 (0%) | 1 (0.2%) | 2 (0%) | 44 (0.1%) |
| 382-02 | Week 1 | 0 (-) | 0 (-) | 152 (8.4%) | 0 (0%) | 0 (-) | 0 (0%) | 0 (0%) |
| 382-02 | Week 2 | 1 (1.5%) | 1 (5.3%) | 0 (-) | 0 (-) | 0 (-) | 0 (0%) | 1 (0.2%) |
| 382-02 | Week 8 | 1 (0.3%) | 0 (0%) | 0 (-) | 0 (-) | 0 (-) | 0 (0%) | 4 (0.1%) |
| 382-02 | Combined | 2 (0.5%) | 1 (1.4%) | 152 (8.4%) | 0 (0%) | 0 (-) | 0 (0%) | 5 (0.1%) |
| 382-07 | Week 1 | 0 (-) | 0 (-) | 192 (13.4%) | 0 (-) | 0 (0%) | 0 (0%) | 0 (0%) |
| 382-07 | Week 2 | 22 (27.2%) | 15 (22.1%) | 0 (-) | 0 (-) | 0 (0%) | 0 (0%) | 1 (0.1%) |
| 382-07 | Week 8 | 3 (0.7%) | 0 (0%) | 0 (-) | 0 (-) | 0 (-) | 0 (0%) | 3 (0.1%) |
| 382-07 | Combined | 25 (4.9%) | 15 (10.9%) | 192 (13.4%) | 0 (-) | 0 (0%) | 0 (0%) | 4 (0.1%) |
| 382-08 | Post-2nd dose<br>week 26 | 0 (-) | 0 (-) | 0 (-) | 0 (0%) | 2 (0.7%) | 0 (0%) | 74 (0.3%) |
| 382-08 | Week 1 | 0 (-) | 0 (-) | 184 (2.1%) | 0 (0%) | 0 (-) | 0 (-) | 0 (0%) |
| 382-08 | Week 2 | 35 (22.3%) | 8 (10.7%) | 0 (-) | 0 (-) | 0 (0%) | 0 (0%) | 2 (0.2%) |
| 382-08 | Week 8 | 48 (29.8%) | 10 (17.5%) | 0 (-) | 0 (-) | 0 (-) | 0 (0%) | 0 (0%) |
| 382-08 | Combined | 83 (26.1%) | 18 (13.6%) | 184 (2.1%) | 0 (0%) | 2 (0.7%) | 0 (0%) | 76 (0.3%) |
| 382-13 | Post-2nd dose<br>week 26 | 0 (-) | 0 (-) | 0 (0%) | 0 (0%) | 0 (0%) | 0 (0%) | 31 (0.2%) |
| 382-13 | Week 1 | 0 (-) | 0 (-) | 270 (9.9%) | 0 (0%) | 0 (-) | 0 (0%) | 0 (0%) |
| 382-13 | Week 2 | 157 (17.2%) | 75 (22.1%) | 0 (-) | 0 (-) | 0 (-) | 0 (0%) | 0 (0%) |
| 382-13 | Week 8 | 192 (22%) | 69 (59.5%) | 0 (-) | 0 (-) | 0 (-) | 0 (0%) | 3 (0.1%) |
| 382-13 | Combined | 349 (19.5%) | 144 (31.6%) | 270 (9.9%) | 0 (0%) | 0 (0%) | 0 (0%) | 34 (0.2%) |
| 382-15 | Week 1 | 0 (-) | 0 (-) | 27 (4.8%) | 0 (0%) | 0 (-) | 0 (0%) | 0 (0%) |
| 382-15 | Week 2 | 403 (33%) | 33 (19.1%) | 0 (-) | 0 (-) | 0 (-) | 3 (0.1%) | 1 (0%) |
| 382-15 | Week 8 | 370 (37.4%) | 131 (44%) | 0 (-) | 0 (-) | 0 (0%) | 2 (0.1%) | 5 (0.3%) |
| 382-15 | Combined | 773 (34.9%) | 164 (34.8%) | 27 (4.8%) | 0 (0%) | 0 (0%) | 5 (0.1%) | 6 (0.1%) |
| Combined |  | 1672<br>(21.8%) | 488 (27.7%) | 1309 (7.3%) | 0 (0%) | 3 (0.4%) | 7 (0%) | 169<br>(0.2%) |

## References

1. Krause, P. R. et al. SARS-CoV-2 Variants and Vaccines. N. Engl. J. Med. 385, 179–186 (2021).

2. Turner, J. S. et al. SARS-CoV-2 mRNA vaccines induce persistent human germinal centre responses. Nature 596, 109–113 (2021).

3. Laidlaw, B. J. & Ellebedy, A. H. The germinal centre B cell response to SARS-CoV-2. Nat. Rev. Immunol. 22, 7–18 (2022).

4. Amanat, F. et al. SARS-CoV-2 mRNA vaccination induces functionally diverse antibodies to NTD, RBD and S2. Cell S0092867421007066 (2021) doi:10.1016/j.cell.2021.06.005.

5. Muecksch, F. et al. Increased memory B cell potency and breadth after a SARS-CoV-2 mRNA boost. Nature 607, 128–134 (2022).

6. Goel, R. R. et al. Efficient recall of Omicron-reactive B cell memory after a third dose of SARS-CoV-2 mRNA vaccine. Cell 185, 1875–1887.e8 (2022).

7. Rodda, L. B. et al. Imprinted SARS-CoV-2-specific memory lymphocytes define hybrid immunity. Cell 185, 1588–1601.e14 (2022).

8. Pérez-Then, E. et al. Neutralizing antibodies against the SARS-CoV-2 Delta and Omicron variants following heterologous CoronaVac plus BNT162b2 booster vaccination. Nat. Med. 28, 481–485 (2022).

9. Sette, A. & Crotty, S. Immunological memory to SARS-CoV-2 infection and COVID-19 vaccines. Immunol. Rev. imr.13089 (2022) doi:10.1111/imr.13089.

10. Lucas, C. et al. Impact of circulating SARS-CoV-2 variants on mRNA vaccine-induced immunity. Nature 600, 523–529 (2021).

11. Wang, Z. et al. mRNA vaccine-elicited antibodies to SARS-CoV-2 and circulating variants. Nature (2021) doi:10.1038/s41586-021-03324-6.

12. Andrews, N. et al. Covid-19 Vaccine Effectiveness against the Omicron (B.1.1.529) Variant. N. Engl. J. Med. 386, 1532–1546 (2022).

13. Kuhlmann, C. et al. Breakthrough infections with SARS-CoV-2 omicron despite mRNA vaccine booster dose. The Lancet 399, 625–626 (2022).

14. Schmidt, F. et al. Plasma neutralization of the SARS-CoV-2 omicron variant. N. Engl. J. Med. 386, 599–601 (2022).

15. Cameroni, E. et al. Broadly neutralizing antibodies overcome SARS-CoV-2 Omicron antigenic shift. Nature 602, 664–670 (2022).

16. Cele, S. et al. Omicron extensively but incompletely escapes Pfizer BNT162b2 neutralization. Nature 602, 654–656 (2022).

17. Falsey, A. R. et al. SARS-CoV-2 Neutralization with BNT162b2 Vaccine Dose 3. N. Engl. J. Med. 385, 1627– 1629 (2021).

18. Bowen, J. E. et al. Omicron spike function and neutralizing activity elicited by a comprehensive panel of vaccines. Science eabq0203 (2022) doi:10.1126/science.abq0203.

19. Muik, A. et al. Neutralization of SARS-CoV-2 Omicron by BNT162b2 mRNA vaccine–elicited human sera. Science 375, 678–680 (2022).

20. Chalkias, S., et al. Safety, Immunogenicity and Antibody Persistence of a Bivalent Beta-Containing Booster Vaccine. https://www.researchsquare.com/article/rs-1555201/v1 (2022) doi:10.21203/rs.3.rs-1555201/v1.

21. Scheaffer, S. M. et al. Bivalent SARS-CoV-2 mRNA vaccines increase breadth of neutralization and protect against the BA.5 Omicron variant. http://biorxiv.org/lookup/doi/10.1101/2022.09.12.507614(2022) doi:10.1101/2022.09.12.507614.

22. Purtha, W. E., Tedder, T. F., Johnson, S., Bhattacharya, D. & Diamond, M. S. Memory B cells, but not long- lived plasma cells, possess antigen specificities for viral escape mutants. J. Exp. Med. 208, 2599–2606 (2011).

23. Lederer, K. et al. Germinal center responses to SARS-CoV-2 mRNA vaccines in healthy and immunocompromised individuals. Cell S0092867422001386 (2022) doi:10.1016/j.cell.2022.01.027.

24. Röltgen, K. et al. Immune imprinting, breadth of variant recognition, and germinal center response in human SARS-CoV-2 infection and vaccination. Cell 185, 1025–1040.e14 (2022).

25. Kim, W. et al. Germinal centre-driven maturation of B cell response to mRNA vaccination. Nature 604, 141– 145 (2022).

26. Case, J. B. et al. Neutralizing antibody and soluble ACE2 inhibition of a replication-competent VSV-SARS- CoV-2 and a clinical isolate of SARS-CoV-2. Cell Host Microbe 28, 475–485.e5 (2020).

27. Case, J. B. et al. Neutralizing antibody and soluble ACE2 inhibition of a replication-competent VSV-SARS- CoV-2 and a clinical isolate of SARS-CoV-2. Cell Host Microbe 28, 475–485.e5 (2020).

28. Liu, Z. et al. Identification of SARS-CoV-2 spike mutations that attenuate monoclonal and serum antibody neutralization. Cell Host Microbe 29, 477–488.e4 (2021).

29. Turner, J. S. et al. Human germinal centres engage memory and naive B cells after influenza vaccination. Nature 586, 127–132 (2020).

30. Ellebedy, A. H. et al. Defining antigen-specific plasmablast and memory B cell subsets in human blood after viral infection or vaccination. Nat. Immunol. 17, 1226–1234 (2016).

31. Radbruch, A. et al. Competence and competition: the challenge of becoming a long-lived plasma cell. Nat. Rev. Immunol. 6, 741–750 (2006).

32. Weisel, F. J., Zuccarino-Catania, G. V., Chikina, M. & Shlomchik, M. J. A temporal switch in the germinal center determines differential output of memory B and plasma cells. Immunity 44, 116–130 (2016).

33. Ellebedy, A. H. et al. Adjuvanted H5N1 influenza vaccine enhances both cross-reactive memory B cell and strain-specific naive B cell responses in humans. Proc. Natl. Acad. Sci. 117, 17957–17964 (2020).

34. Zost, S. J. et al. Potently neutralizing and protective human antibodies against SARS-CoV-2. Nature 584, 443– 449 (2020).

35. Chen, P. et al. SARS-CoV-2 neutralizing antibody LY-CoV555 in outpatients with COVID-19. N. Engl. J. Med. 384, 229–237 (2021).

36. Hansen, J. et al. Studies in humanized mice and convalescent humans yield a SARS-CoV-2 antibody cocktail. Science 369, 1010–1014 (2020).

37. Alsoussi, W. B. et al. A potently neutralizing antibody protects mice against SARS-CoV-2 infection. J. Immunol. 205, 915–922 (2020).

38. Chen, R. E. et al. In vivo monoclonal antibody efficacy against SARS-CoV-2 variant strains. Nature (2021) doi:10.1038/s41586-021-03720-y.

39. Francis, T. On the Doctrine of Original Antigenic Sin. Proc. Am. Philos. Soc. 104, 572–578 (1953).

40. Gostic, K. M., Ambrose, M., Worobey, M. & Lloyd-Smith, J. O. Potent protection against H5N1 and H7N9 influenza via childhood hemagglutinin imprinting. Science 354, 722–726 (2016).

41. . Zang, R., et al. TMPRSS2 and TMPRSS4 promote SARS-CoV-2 infection of human small intestinal enterocytes. Sci. Immunol. 5, eabc3582 (2020).

42. Stadlbauer, D. et al. SARS-CoV-2 seroconversion in humans: a detailed protocol for a serological assay, antigen production, and test setup. Curr. Protoc. Microbiol. 57, (2020).

43. Fairhead, M. & Howarth, M. Site-specific biotinylation of purified proteins using BirA. Methods Mol. Biol. 1266, 171–184 (2015).

44. Chen, R. E. et al. Resistance of SARS-CoV-2 variants to neutralization by monoclonal and serum-derived polyclonal antibodies. Nat. Med. (2021) doi:10.1038/s41591-021-01294-w.

45. Liu, Z. et al. Identification of SARS-CoV-2 spike mutations that attenuate monoclonal and serum antibody neutralization. Cell Host Microbe 29, 477–488.e4 (2021).

46. VanBlargan, L. A. et al. A potently neutralizing SARS-CoV-2 antibody inhibits variants of concern by utilizing unique binding residues in a highly conserved epitope. Immunity 54, 2399–2416.e6 (2021).

47. Mudd, P. A. et al. SARS-CoV-2 mRNA vaccination elicits a robust and persistent T follicular helper cell response in humans. Cell S0092867421014896 (2021) doi:10.1016/j.cell.2021.12.026.

48. Wrammert, J. et al. Broadly cross-reactive antibodies dominate the human B cell response against 2009 pandemic H1N1 influenza virus infection. J. Exp. Med. 208, 181–193 (2011).

49. Smith, K. et al. Rapid generation of fully human monoclonal antibodies specific to a vaccinating antigen. Nat. Protoc. 4, 372–384 (2009).

50. Wrammert, J. et al. Rapid cloning of high-affinity human monoclonal antibodies against influenza virus. Nature 453, 667–671 (2008).

51. Nachbagauer, R. et al. Broadly Reactive Human Monoclonal Antibodies Elicited following Pandemic H1N1 Influenza Virus Exposure Protect Mice against Highly Pathogenic H5N1 Challenge. J. Virol. 92, 1–17 (2018).

52. Brochet, X., Lefranc, M.-P. & Giudicelli, V. IMGT/V-QUEST: the highly customized and integrated system for IG and TR standardized V-J and V-D-J sequence analysis. Nucleic Acids Res. 36, W503–W508 (2008).

53. Giudicelli, V., Brochet, X. & Lefranc, M.-P. IMGT/V-QUEST: IMGT Standardized Analysis of the Immunoglobulin (IG) and T Cell Receptor (TR) Nucleotide Sequences. Cold Spring Harb. Protoc. 2011, pdb.prot5633-pdb.prot5633 (2011).

54. Vander Heiden, J. A., et al. pRESTO: a toolkit for processing high-throughput sequencing raw reads of lymphocyte receptor repertoires. Bioinformatics 30, 1930–1932 (2014).

55. Turner, J. S. et al. SARS-CoV-2 mRNA vaccines induce persistent human germinal centre responses. Nature 596, 109–113 (2021).

56. Schmitz, A. J. et al. A vaccine-induced public antibody protects against SARS-CoV-2 and emerging variants. Immunity 54, 2159–2166.e6 (2021).

57. Ye, J., Ma, N., Madden, T. L. & Ostell, J. M. IgBLAST: an immunoglobulin variable domain sequence analysis tool. Nucleic Acids Res. 41, W34–W40 (2013).

58. Gadala-Maria, D., Yaari, G., Uduman, M. & Kleinstein, S. H. Automated analysis of high-throughput B-cell sequencing data reveals a high frequency of novel immunoglobulin V gene segment alleles. Proc. Natl. Acad. Sci. 112, E862–E870 (2015).

59. Zhou, J. Q. & Kleinstein, S. H. Cutting Edge: Ig H Chains Are Sufficient to Determine Most B Cell Clonal Relationships. J. Immunol. 203, 1687–1692 (2019).

60. Gupta, N. T. et al. Hierarchical Clustering Can Identify B Cell Clones with High Confidence in Ig Repertoire Sequencing Data. J. Immunol. 198, 2489–2499 (2017).

61. Gupta, N. T. et al. Change-O: a toolkit for analyzing large-scale B cell immunoglobulin repertoire sequencing data. Bioinformatics 31, 3356–3358 (2015).

62. Gu, Z., Gu, L., Eils, R., Schlesner, M. & Brors, B. circlize implements and enhances circular visualization in R. Bioinformatics 30, 2811–2812 (2014).

63. Wolf, F. A., Angerer, P. & Theis, F. J. SCANPY: large-scale single-cell gene expression data analysis. Genome Biol. 19, 15 (2018).

64. Turner, J. S., et al. Germinal centres foster recall and de novo human B cell responses to influenza vaccination. (2020).

65. Haebe, S. et al. Single-cell analysis can define distinct evolution of tumor sites in follicular lymphoma. Blood 137, 2869–2880 (2021).

66. Mourcin, F. et al. Follicular lymphoma triggers phenotypic and functional remodeling of the human lymphoid stromal cell landscape. Immunity 54, 1788–1806.e7 (2021).

